# Foraging in conceptual spaces: hippocampal oscillatory dynamics underlying searching for concepts in memory

**DOI:** 10.1101/2025.10.07.680852

**Authors:** Simone Viganò, Giuliano Giari, Roberto Mai, Christian F. Doeller, Roberto Bottini

## Abstract

How does the brain access stored knowledge? It has been proposed that conceptual search engages neurocognitive processes similar to foraging in physical space. We tested this idea using intracranial EEG in patients performing a verbal fluency task, where they spontaneously explored their own knowledge of the world, sampling words from semantic memory. We found that hippocampal theta power increased during conceptual search and scaled with the semantic distance between successive words, paralleling dynamics observed in spatial navigation. Critically, people transitioned between conceptual clusters, resembling transitions between resource patches in foraging behavior. These shifts were marked by enhanced theta-gamma coupling, both within the hippocampus and between the hippocampus and lateral temporal cortex, a key hub of the semantic network. These findings support a mechanistic account of memory search grounded in navigation and foraging principles, suggesting that the hippocampus orchestrates local computations and long-range interactions to enable flexible retrieval of conceptual knowledge.

## Introduction

The ability to search for resources is fundamental to survival, manifesting across animal species in behaviors such as foraging or hunting. Humans extend this capacity into the domain of abstract, non-spatial knowledge, for instance browsing for information on the web or engaging in more “internal” forms of search, like retrieving ideas or concepts from memory. Philosophers and cognitive scientists have long proposed that such mental foraging may rely on mechanisms analogous to those used in the navigation and exploration of physical spaces (St. Augustine, Confessions, Book X, 398; James, 1890; Lakoff & Johnson, 1999; Gärdenfors, 2000; Todd & Hills 2020; Jaynes 1978). In semantic fluency tasks, for instance, where individuals have to recollect items belonging to a conceptual category stored in memory (e.g., “Tell me all the animals that come to your mind”), responses tend to cluster by similarity in their meaning before shifting to new subcategories (e.g., listing farm animals before switching to birds; Hills et al., 2012). This parallels “area-restricted” search patterns seen in foraging animals, that balance global exploration of the environment with local exploitation of specific locations (Laing, 1937; Tinbergen et al., 1967; Hills et al., 2015; see reviews by Todd & Hills, 2020; Dorfman et al., 2022). Consistently with cognitive science intuitions, neuroscientific studies have suggested that the brain may repurpose spatial coding systems to structure abstract knowledge. For instance, activity in the hippocampal-entorhinal system, a brain region central to spatial navigation and episodic memory (Tolman, 1949; Moser et al., 2017; Squire, 2004; Dickerson & Eichenbaum, 2010; Burgess et al. 2002), has been implicated in the organization of conceptual spaces and relational memory of non-spatial information (Constantinescu et al., 2016; Theves et al., 2019, 2020; Viganò & Piazza, 2020; Viganò et al., 2021, 2023; Park et al., 2021; Barnaveli et al., 2024; Qasim et al., 2023; Jasmin et al. 2025; Aronov et al. 2017; see also Bellmund et al., 2018; Behrens et al., 2018; Buzsáki & Moser, 2013; Bottini & Doeller, 2020). More recent evidence further links hippocampal activity to performance in semantic fluency tasks (Gleissner & Elger, 2001; Sheldon & Moscovitch, 2012; Glikmann-Johnston et al., 2015; Lee et al., 2021; Lundin et al., 2023; Nour et al., 2023), suggesting that its role may extend beyond spatial and episodic memory into semantic cognition. Yet, the neural mechanisms underlying internal search remain poorly understood.

Here, we address this gap by leveraging intracranial electrophysiology for its high temporal resolution and anatomical specificity. This allowed us to directly record neural activity from the human hippocampus (Figure 1a) during semantic foraging, while people spontaneously sampled concepts from their semantic knowledge of the world (Figure 1b). In particular, we focused on oscillatory mechanisms, because the rhythmic neural activity in the hippocampus is thought to structure internal representations across both space and time, organizing cell assemblies into coherent sequences that support memory-guided behavior (Buzsáki, 2005; Buzsáki & Moser, 2013). In the spatial domain, hippocampal theta oscillations have indeed been shown to support spatial navigation and memory (e.g., Bush et al. 2017; Winson et al. 1978; Seger et al. 2023), and thus we asked whether and how they might provide a temporal scaffold for simulating and traversing relational structures more generally.

**Figure 1.**
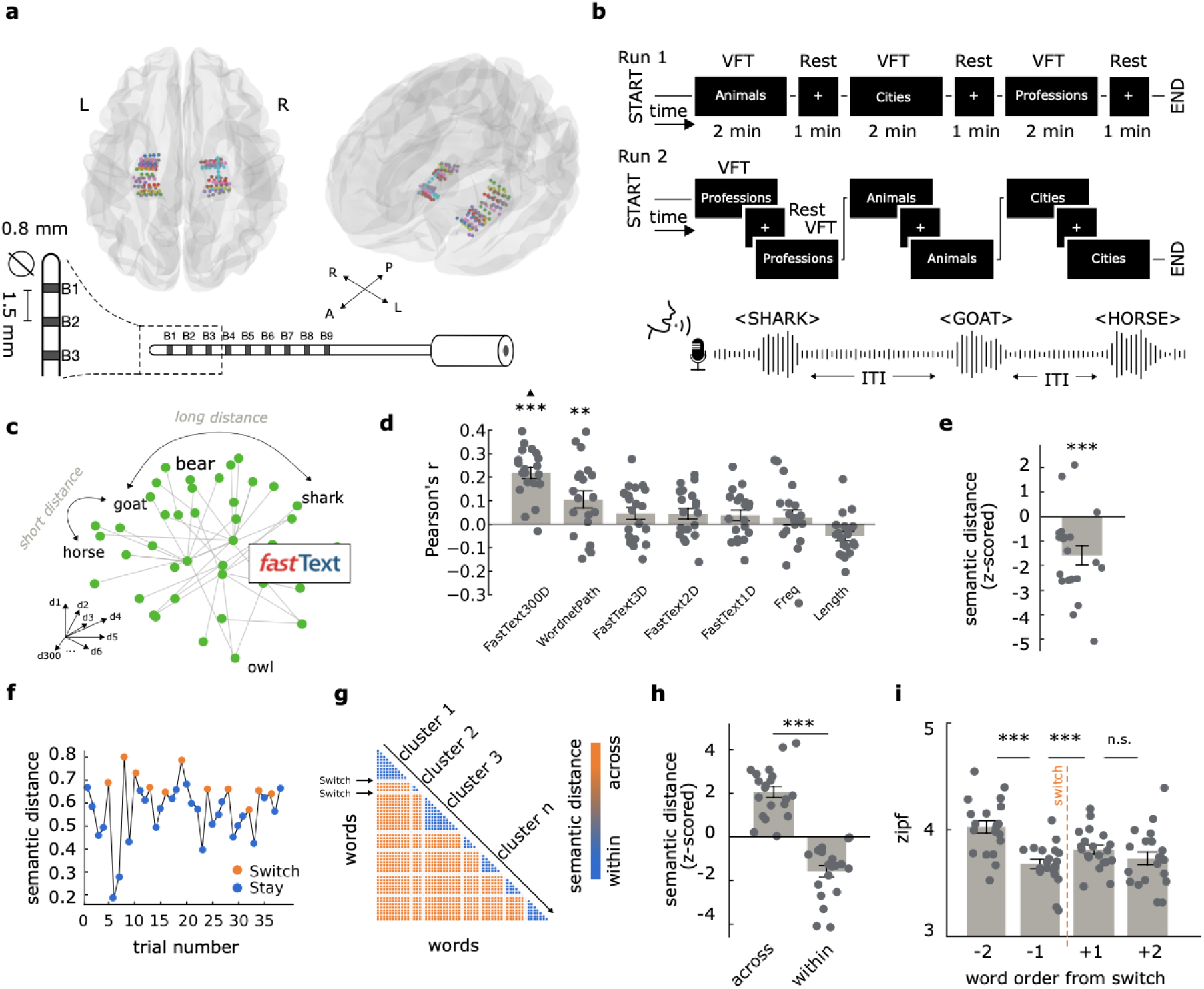
Experimental design and behavioral metrics of internal semantic search. **(a)** Stereotactic intracranial EEG (sEEG) recordings were acquired from 20 patients undergoing epilepsy monitoring, with electrodes targeting bilateral hippocampi. The glass brain illustrates electrode contact locations in MNI space, color-coded by participants. Below, a schematic of a representative depth electrode is shown, with labeled contacts (e.g., B1, B2, B3). **(b)** Schematic of the categorical Verbal Fluency Task (cVFT), in which participants generated as many animal, or cities, or profession names as possible within 120-second blocks. In Run 1, foraging blocks of different categories were interspersed with 60-s rest periods, whereas in Run 2 the rest periods were embedded within two foraging blocks of the same category. Category order was randomized across participants. **(c)** Each spoken word was modeled as a high-dimensional vector embedding (FastText, 300 dimensions). Semantic distance between words was computed as 1 minus the cosine similarity between successive vectors. **(d)** Fisher-transformed Pearson’s correlations between inter-word time intervals (ITI) and the semantic or linguistic distance between successive words. Distances include FastText-based semantic distance (300D and low-dimensional PCA projections), WordNet path distance, absolute differences in lexical frequency (Freq), and word length (Length). High dimensional semantic vectors (Fasttext300D) were the best predictor of inter-word latency (▴ p <.05 compared to every other model). **(e)** Semantic distances (FastText300D) between successive words, normalized (z-scored) against 1,000 trial-wise permutations. **(f)** Example trial showing how word-to-word semantic distance is used to classify transitions as *switches* (across clusters) or *stays* (within cluster), following Lundin et al. 2023. **(g-h)** Semantic distances between words produced *within* vs *across* clusters **(i)** Linguistic frequency of generated words plotted as a function of their order relative to switch transitions ***p < .001, **p < .005, *p < .01. Dots represent individual participants. Error bars represent standard error of the mean.

## Results

### The free internal exploration of personal conceptual spaces mimics spatial foraging behavior

To investigate the behavioral and neural dynamics of semantic search, we asked twenty patients with drug-resistant epilepsy undergoing stereo-electroencephalography (sEEG; Figure 1a) to perform a categorical verbal fluency task (Bousfield & Sedgewick, 1944). Participants were instructed to name as many items as possible from a given semantic category (animals, cities, or professions) across multiple runs and blocks (Figure 1b). On average, participants generated 17.01 words (SD = 5.89) per block, with a mean inter-word interval (ITI) of 6.79 s (SD = 2.23). Word counts and ITIs did not differ significantly across categories (Word count: F(2,56) = 0.94, p = 0.396; ITI: F(2,56) = 0.277, p = 0.758) or repetition (Word count: F(2,57) = 0.536, p = 0.587; ITI: F(2,57) = 0.695, p = 0.503).

We modeled each verbalized word as a vector in a 300-dimensional semantic embedding space (Bojanowski et al., 2016), with semantic distance operationalized as the inverse of cosine similarity (Mikolov et al. 2013; Piantadosi et al., 2024)(Figure 1c; see Methods). This measure of dissimilarity was the best predictor, out of seven that we considered (see Methods), of participants’ inter-word latency (t(19)=9.118, p < 0.001, all pairwise p < .05; Figure 1d). This suggests that internal search is sensitive to conceptual proximity, and might mirror physical navigation, where greater distances (in our case, dissimilarities) take longer to traverse (at a constant speed). Moreover, participants’ word sequences showed a strong tendency toward local semantic proximity: consecutively produced words were significantly closer in semantic space than expected by chance (t(19) = −4.017, p < 0.001, M = −1.572, SD = 1.75, CI = [−2.52, −0.62]; Figure 1e; see Methods). Occasionally however, participants deviated from exploiting local semantic regions, producing words at greater semantic distances (and thus more dissimilar) before returning to tighter clusters (Figure 1f)(see Methods). These fewer transitions, occurring on average 33% of the time, were interpreted as semantic switches to new sub-clusters of related items (e.g., from “farm animals” to “birds”; similar to the occasional long-range global exploration observed in animals during foraging, see Hills et al., 2012; Lundin et al., 2023; Kumar et al., 2024). Supporting this interpretation, semantic distances were significantly lower within clusters (between two switches) than across clusters (spanning at least one switch)(t(19) = −7.066, p < 0.001; Figure 1g-h). Semantic distances within clusters were still statistically significantly correlated with inter-word latency (t(19)=8.577, p<0.001). Overall, these behavioral dynamics, consistent with principles of area-restricted search and optimal foraging theory (Hills et al., 2012; 2015; Todd & Hills, 2020), suggest that participants alternated between phases of local exploitation and broader exploration of their semantic memory. Previous studies have suggested that transitions to new clusters might co-occur with changes in linguistic frequency of sampled concepts (Hills et al. 2015). We observed that cluster switches were preceded by a significant drop in linguistic frequency at cluster boundaries (F(3,76)=6.067, p<0.001; −2 to −1: t(19)=-3.460, p=0.002; −1 to +1: t(19)=3.066, p=0.006; +1 to +2: t(19)=-1.2, p=0.241; Figure 1i), possibly reflecting the depletion of available local resources (that is, easily accessible, frequent words in the current semantic cluster) prior to the switch.

Together, these findings indicate that semantic foraging is influenced by at least four factors:

(i) semantic distance between concepts, (ii) retrieval latency, (iii) balance between local exploitation and global exploration via semantic switches, and (iv) word frequency. We then used these behavioral metrics as a framework for studying the neural correlates of semantic search in the hippocampus.

### Searching and finding concepts is accompanied by frequency-specific changes in hippocampal oscillatory power

All participants had depth electrodes targeting the medial temporal lobe, including the hippocampus (174 bipolar contacts in total; Supplementary Table 1; Figure 1a), enabling us to record local field potentials (LFPs) during semantic foraging. Guided by prior work suggesting that memory-related neural signatures emerge prior to verbalization (Addante et al., 2011; Fell et al., 2011; Solomon et al., 2019), we conducted a time-frequency analysis in the 1-second window preceding word onset (−1 to 0 s) and controlled for the aperiodic component of the spectrum using IRASA (Wen & Liu, 2016). We then contrasted the oscillatory time-frequency spectrum with the subsequent 1s post-onset interval (Figure 2a)(see Methods).

**Figure 2.**
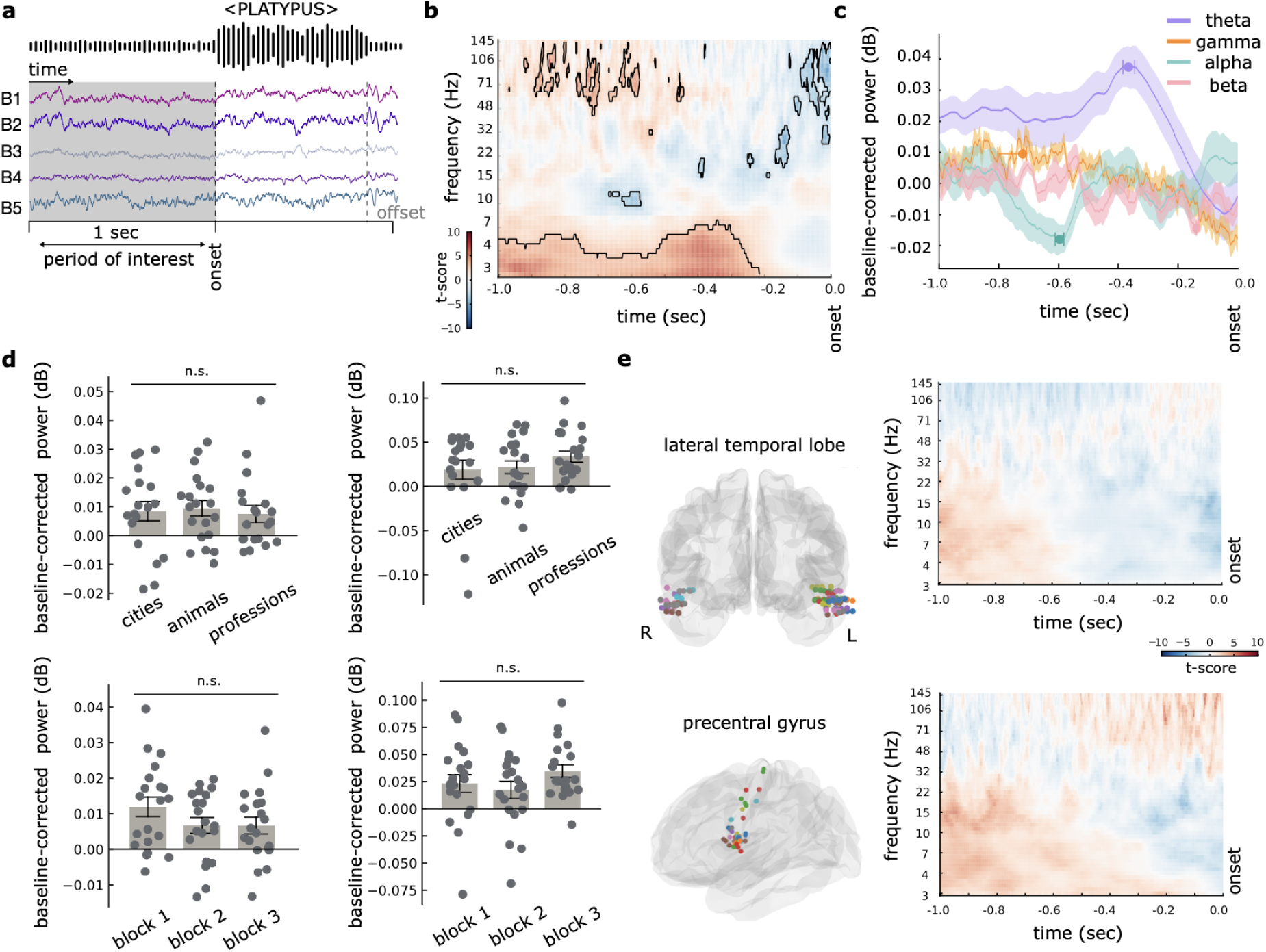
Hippocampal time-frequency dynamics during internal semantic search. **(a)** Analysis logic. We focused on the 1-second window preceding word onset (“pre-onset”), corresponding to internal semantic search and conceptual retrieval, and compared it to a 1-second post-onset baseline, reflecting articulation processes. Control analyses confirmed that the observed effects reported below are robust to baseline choice (see Supplementary Figure 1a). **(b)** Time–frequency representation of hippocampal activity during the pre-onset interval, showing increased theta (3-7.2 Hz) and gamma (>41 Hz) power, and decreased alpha and beta power. Black contours indicate significant clusters (p < 0.05, FDR-corrected). **(c)** Time-resolved visualization of power changes across canonical frequency bands. Shaded area represents standard error of the mean. Dots mark peak changes; error bars represent 95% bootstrapped CI. **(d)** No statistically significant effect of category or block were observed for theta (left) or gamma (right) power across conceptual categories (top) or blocks (bottom). Dots represent individual participants. Error bars represent standard error of the mean. **(e)** Control analyses in non-hippocampal regions (lateral temporal cortex (top) and precentral gyrus (bottom)) did not yield any statistically significant clusters at p < .05 FDR-corrected, underscoring the anatomical specificity of the reported main effect.

This analysis revealed a robust increase in theta (3-7.2 Hz) and high gamma (>41 Hz) oscillatory power, along with a significant decrease in alpha (9.8–11.5. Hz) and beta (15.8–34.8 Hz) bands (all p < 0.05, FDR-corrected; Figure 2b). Different temporal profiles were observed for different frequency bands. While theta power was elevated almost throughout the entire pre-verbalization window (−1 to −.2 s), peaking at −0.378 s (bootstrapped CI = [-0.384, 0.372] s), the gamma increase peaked earlier (−0.75 s, CI=[-0.866, −0.708] s) and was temporally restricted between −0.968 and −0.444 s. Alpha and beta power decreases were more transient, confined to narrow windows (alpha: −0.668 to −0.572 s, peak at −0.6 s, CI=[-0.611, −0.596] s; beta: −0.398 to −0.124 s, peak at −0.138 s; CI = [-0.138, −0.138] s)(Figure 2c). The effects were consistent across semantic categories (theta: F(2,56) = 0.926, p = 0.401; gamma: F(2,56) = 0.106, p = 0.899; alpha: F(2, 56) = 1.346, p=0.268; beta: F(2, 56) = 0.087, p=0.915) (Figure 2d - top), block repetitions (theta: F(2,57) = 1.40, p = 0.254; gamma: F(2,57) = 1.526, p = 0.226; alpha: F(2, 57) = 1.767, p=0.179; beta: F(2, 57) =0.523, p = 0.595)(Figure 2d - bottom), and alternative baseline periods (theta: F(3,76)=1.431, p=0.240, gamma: F(3,76)=0.717, p=0.544, alpha: F(3, 76)=0.285, p=0.835, beta: F(3, 76)=0.395, p=0.756)(Supplementary Figure 1). No statistically significant effect was observed in two control regions, the lateral temporal lobe and the precentral gyrus (Figure 2e). In short, searching and finding concepts during semantic foraging was accompanied by sustained increases in hippocampal theta, similar to what has been observed during spatial navigation and exploration in physical or virtual reality environments. We also observed increased gamma oscillatory activity in the hippocampus, and more focal decreases in its alpha and beta power. We next asked whether these frequency-specific dynamics related to key behavioral features of semantic search, such as inter-word semantic distance, switching, retrieval latency, or word frequency.

### Hippocampal theta oscillations during semantic foraging are modulated by high-dimensional semantic distance between sampled concepts

To identify which aspects of internal search were tracked by hippocampal activity, we followed prior work in the spatial domain (Stangl et al., 2020; Liu et al., 2023) and modeled oscillatory power in each frequency band (theta, gamma, alpha, and beta) using linear mixed-effects models at single trial level (see Methods).

Similarly to what has been observed for travelled distance during spatial navigation in virtual reality environments (Bush et al. 2017), theta power was significantly modulated by semantic distance in high-dimensional FastText space (β = 0.012, 94% HDI = [0.005, 0.0019]), with larger distances predicting stronger theta activity (Figure 3a-b). Interestingly, this effect was statistically absent when modelling semantic distances in a low-dimensional (1D, 2D, or 3D) projection of FastText space (Supplementary Figure 2). We did not observe any statistically significant modulation of gamma, alpha, and beta power (Figure 3c). All these results were robust across alternative analytical approaches (Supplementary Figure 3). Interestingly, the modulation of theta power as a function of semantic distance was replicated within the hippocampus also when considering only trials within clusters (β = 0.01, 94% HDI = [0.001, 0.019]), indicating that this effect was not driven by switch trials (Supplementary Figure 4).

**Figure 3.**
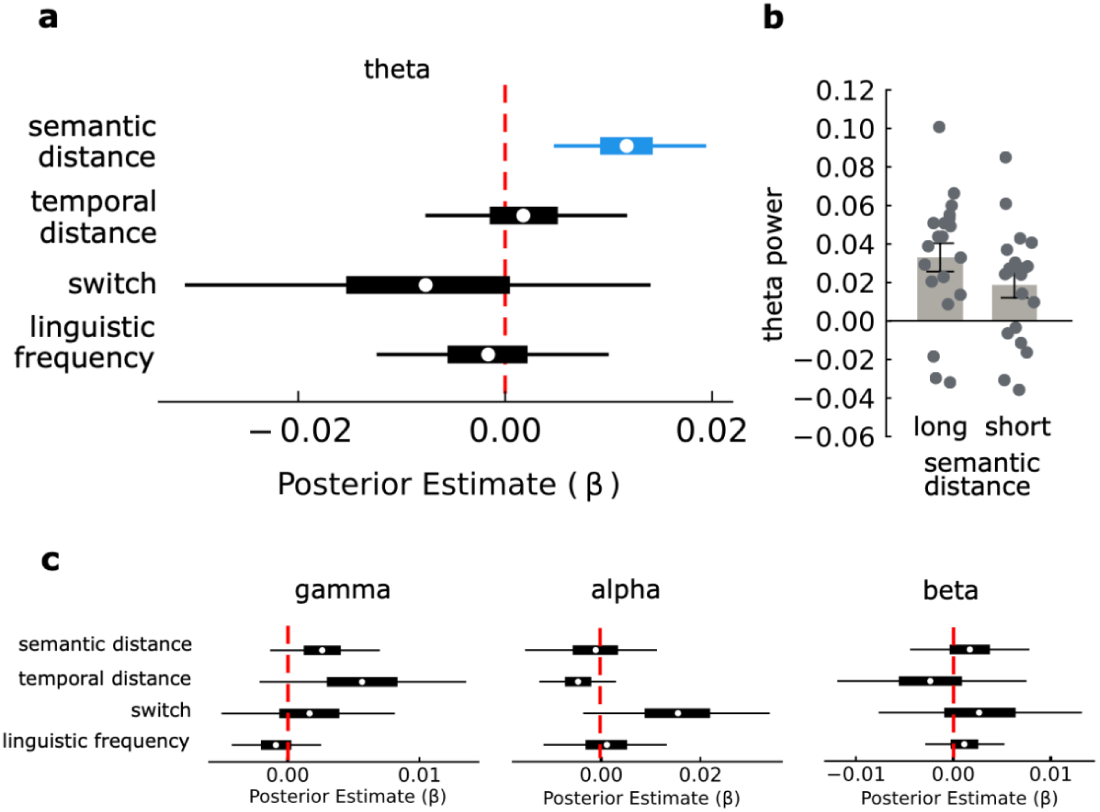
Hippocampal theta power tracks semantic distance during internal search. **(a)** Posterior estimates from a Bayesian linear mixed-effects model predicting hippocampal theta power during a verbal fluency task. White circles indicate posterior medians; thick bars represent the central 50% credible interval (inter-quartile range); thin lines denote 94% high-density intervals. A selective effect of semantic distance emerged (blue), with its 94% interval excluding zero, indicating a reliable increase in theta power with greater conceptual distance between successive words. Other regressors (temporal distance (β = 0.002, 94% HDI = [−0.008, 0.012]), cluster switch (β = −0.007, 94% HDI = [−0.031, 0.014]), and lexical frequency (β = −0.002, 94% HDI = [−0.012, 0.01])) showed no credible effects. The red dashed line marks the null value (β = 0). **(b)** Visualization of theta power for long vs short semantic distances (median split within participant). Dots represent individual participants. Error bars represent standard error of the mean. **(c)** Same as a, but for the other frequency bands: gamma (left): semantic distance: (β = 0.003, 94% HDI = [−0.001, 0.007]), temporal distance (β = 0.006, 94% HDI = [−0.002, 0.014]), cluster switch (β = 0.002, 94% HDI = [−0.005, 0.008]), and lexical frequency (β = −0.001, 94% HDI = [−0.004, 0.003]);, alpha (center): semantic distance: (β = −0.001, 94% HDI = [−0.015, 0.011]), temporal distance (β = −0.005, 94% HDI = [−0.013, 0.003]), cluster switch (β = 0.015, 94% HDI = [−0.004, 0.034]), and lexical frequency (β = 0.001, 94% HDI = [−0.012, 0.013]); beta(right): semantic distance: (β = 0.002, 94% HDI = [−0.004, 0.008]), temporal distance (β = −0.002, 94% HDI = [−0.012, 0.008]), cluster switch (β = 0.003, 94% HDI = [−0.008, 0.013]), and lexical frequency (β = 0.001, 94% HDI = [−0.003, 0.005]).

### Hippocampal gamma oscillations are modulated as a function of theta phase, and their coupling is higher when switching to a new semantic cluster of related concepts

Given prior evidence that hippocampal gamma activity is modulated by theta phase (Colgin et al. 2009; Colgin & Moser, 2010; Lisman & Jensen, 2013), we next investigated whether theta-gamma phase-amplitude coupling (TG-PAC) occurred during semantic search and, if so, whether it reflected any of the behavioral dynamics identified above. TG-PAC has been indeed proposed as a mechanism for memory encoding and retrieval (Griffiths & Jensen, 2023), enabling the representation of sequential information through gamma bursts nested within theta cycles (Lisman & Jensen, 2013), and has also been linked to working memory and cognitive control in hippocampal–prefrontal networks (Daume et al., 2024).

Following previous studies (Tort et al., 2008; Heusser et al., 2016; Kragel et al., 2019; Daume et al., 2024), we estimated TG-PAC as a function of theta (3 to 7.2 Hz) phase and gamma (41-145 Hz) amplitude. This revealed a statistically significant coupling between high-gamma amplitude (83–98 Hz) and high-theta phase (5.2–7.2 Hz) in the −1 to 0 s interval preceding word onset (p = 0.013, cluster-corrected; Figure 4a left) that persisted also after word onset (p = 0.0152, cluster corrected, Figure 4a right). TG-PAC was not induced by increase in the power of theta or gamma, nor influenced by theta waveform shape (Supplementary Figure 5).

**Figure 4.**
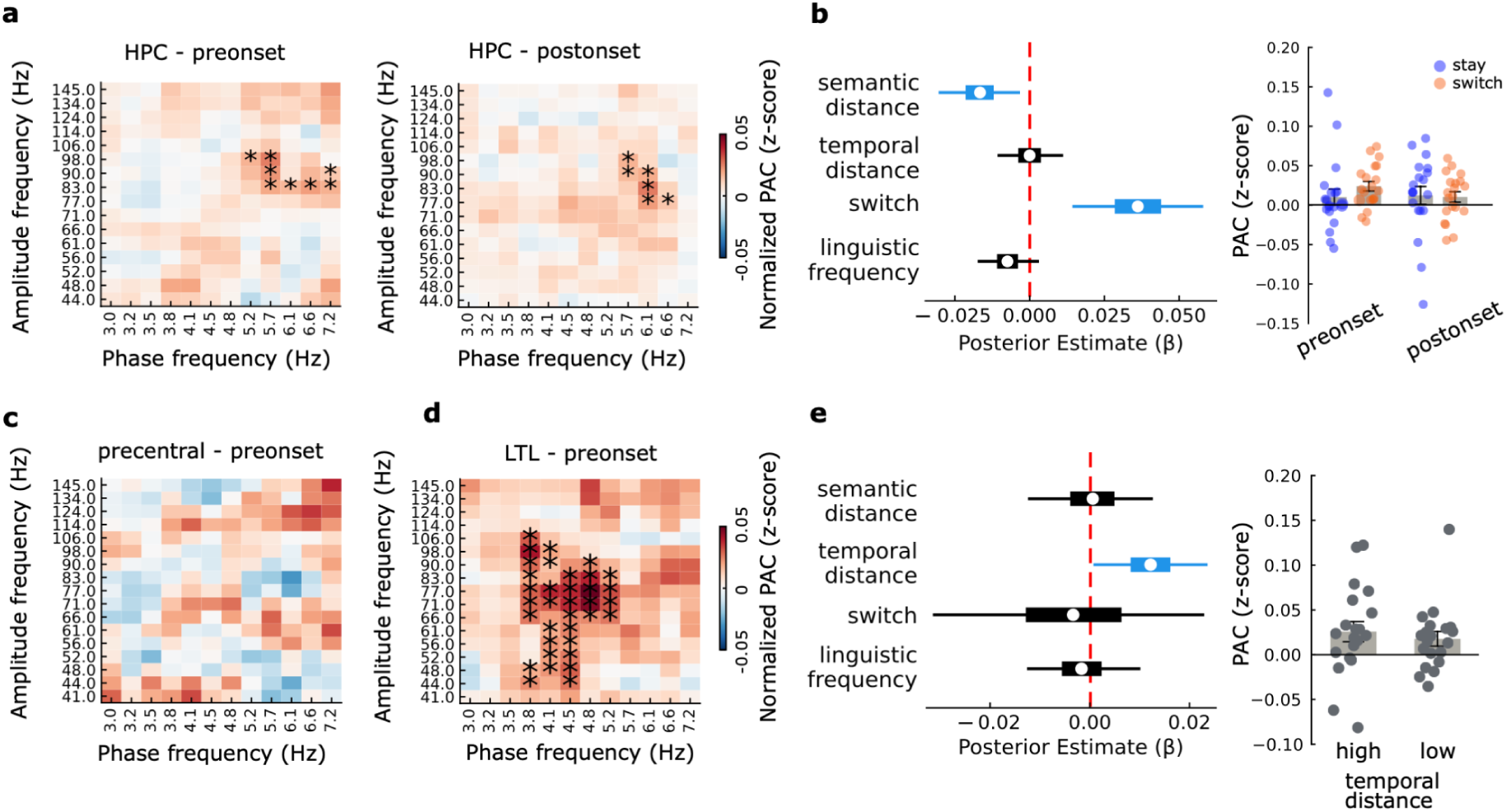
Theta-gamma phase–amplitude coupling (TG-PAC) tracks semantic search behaviour in the hippocampus and lateral temporal cortex. **(a)** Theta–gamma PAC (z-scored) in the hippocampus (HPC) during the 1 s window before (left) and after (right) word production onset. Significant clusters (asterisks) indicate enhanced TG-PAC between theta phase (5.2–7.2 Hz) and gamma amplitude (83–98 Hz) during both epochs (cluster-corrected). **(b)** Left: Bayesian linear mixed-effects model estimates relating hippocampal TG-PAC in the pre-onset window to semantic distance (β = −0.017, 94% HDI = [−0.03, −0.003]), temporal distance (β = 0, 94% HDI = [−0.011, 0.011]), switch (β = 0.036, 94% HDI = [0.014, 0.058]), and lexical frequency (β = −0.007, 94% HDI = [−0.017, 0.003]). Cluster switch showed a robust positive effect (94% high-density interval excludes 0; blue), suggesting that TG-PAC increases during transitions to new sub-clusters. This was not observed in the post-onset window. Similar but weaker effects are observed for semantic distance, with a negative modulation. Right: Visualization of TG-PAC as higher on “switch” trials (orange) than “stay” trials (orange) during pre-onset. Dots represent individual participants. Error bars represent standard error of the mean. (**c–d)** Theta–gamma PAC (z-scored) for precentral cortex and lateral temporal lobe (LTL), respectively. Only LTL exhibited a significant theta–gamma PAC cluster (cluster-corrected). **(e)** Left: Posterior estimates from the same model as (b), applied to LTL PAC. Only temporal distance showed a modest but significant effect (β = 0.012, 94% HDI = [0.001, 0.024]) All other predictors HDIs crossed zero: semantic distance (β = 0, 94% HDI = [−0.012, 0.013]); switch: (β = −0.003, 94% HDI = [−0.031, 0.023]); linguistic frequency (β = −0.002, 94% HDI = [−0.013, 0.01]). Right: Visualization of TG-PAC as higher for larger temporal distances (median split within subject). Dots represent individual participants. Error bars represent standard error of the mean.

We then examined whether TG-PAC tracked specific behavioral features of semantic foraging using the same linear mixed-effects framework we used for analysing modulation of power. Among the predictors tested, TG-PAC was significantly modulated by semantic distance (β = −0.017, 94% HDI = [−0.03, −0.003]) and, crucially, by semantic cluster switching: coupling strength was greater during transitions between clusters than during within-cluster sampling (β = 0.036, 94% HDI = [0.014, 0.058]; Figure 4b). This modulatory effect of switch, but not the one of semantic distance, was mostly robust across different modeling approaches (Supplementary Figure 6), and was not observed after word onset (94% HDI of all parameters crossing zero), with a significant interaction with pre-onset period (β = 0.029, 94% HDI = [0.003, 0.057]) confirming its specific involvement during mental search, rather than verbalization. Interestingly, while we did not observe any statistically significant evidence of TG-PAC in our control region in the precentral gyrus (all p > .36, cluster-corrected, Figure 4c), we did observe statistically significant evidence of coupling in the lateral temporal lobe, during the preonset time window (p = 0.001, cluster-corrected)(Figure 4d)(see Supplementary Figure 7 for controls). In this case however, rather than being modulated by cluster switch as it happened within the hippocampus, TG-PAC within LTL was statistically significantly modulated as a function of the time needed to find the next word, albeit more moderately (β = 0.012, 94% HDI = [0.001, 0.024])(Figure 4e).

Taken together, all these findings suggest that hippocampal theta-gamma coupling may be related to cognitive transitions between distinct regions of semantic space while participants are searching for concepts in their long-term memory. They also indicate that the lateral temporal lobe might serve an additional role in this process. This prompted us to ask whether theta-gamma coupling might act as a bridge between local computations in the hippocampus and distributed cortical processing during semantic exploration.

### Theta-gamma coupling between hippocampus and lateral temporal cortex is enhanced during cluster switches

A major hypothesis on the role of phase-amplitude coupling, indeed, is that it facilitates inter-areal communication by temporally aligning local high-frequency activity with slower rhythms that coordinate long-range synchronization (Buzsáki & Wang 2012; Fries, 2023; Griffiths & Jensen, 2023). While gamma oscillations are typically restricted to local networks due to their fast temporal dynamics and short spatial reach (Ray & Maunsell, 2015; von Stein & Sarnthein, 2000), theta oscillations can synchronize activity across distributed cortical and hippocampal regions (Sirota et al., 2008; Roux et al., 2022). The presence of TG-PAC during our task thus raised the possibility of theta-mediated coordination between the hippocampus and neocortex. The lateral temporal lobe (LTL), in particular, is a key brain area implicated in language and semantic memory (Binder et al., 2009; Fedorenko et al., 2024), and it was conveniently covered by the same shafts targeting the hippocampus, allowing us to test our hypothesis.

We computed cross-regional TG-PAC between hippocampus (HPC) and LTL. We observed statistically significant effects in the period preceding word onset both when considering coupling between phase of hippocampal theta and amplitude of LTL gamma (p = 0.0156, cluster-corrected, Figure 5a top left) as well as when we considered phase of LTL theta and amplitude of hippocampal gamma (p < 0.001, cluster-corrected, Figure 5a top right). The latter effect persisted also after word onset (p < 0.001, cluster-corrected, Figure 5a bottom right). Interestingly, these statistically significant effects involved a lower range of theta (between 3 to 4.8 Hz), differently from the local effects in the hippocampus, which involved a higher one (between 5.2 and 7.2 Hz). Moreover and crucially, cross-regional TG-PAC between hippocampus and LTL was modulated as a function of cluster switch (β = 0.02, 94% HDI = [0.005, 0.036])(Figure 5b): it was stronger when participants were switching to a new region of their semantic space (“exploration”), and weaker when participants were retrieving concepts from the same cluster (“exploitation”) (Figure 5c). This effect seemed to be specific to the interaction between hippocampus and LTL because we did not observe a statistically significant cross-regional TG-PAC between hippocampus and precentral gyrus before word onset, when participants were searching for, and finding, the next word. Moreover, although we did observe a significant coupling between these two regions after word onset, this was not significantly modulated by any of our behavioral predictors (Supplementary Figure 8), potentially linking this effect to articulation and verbalization.

**Figure 5.**
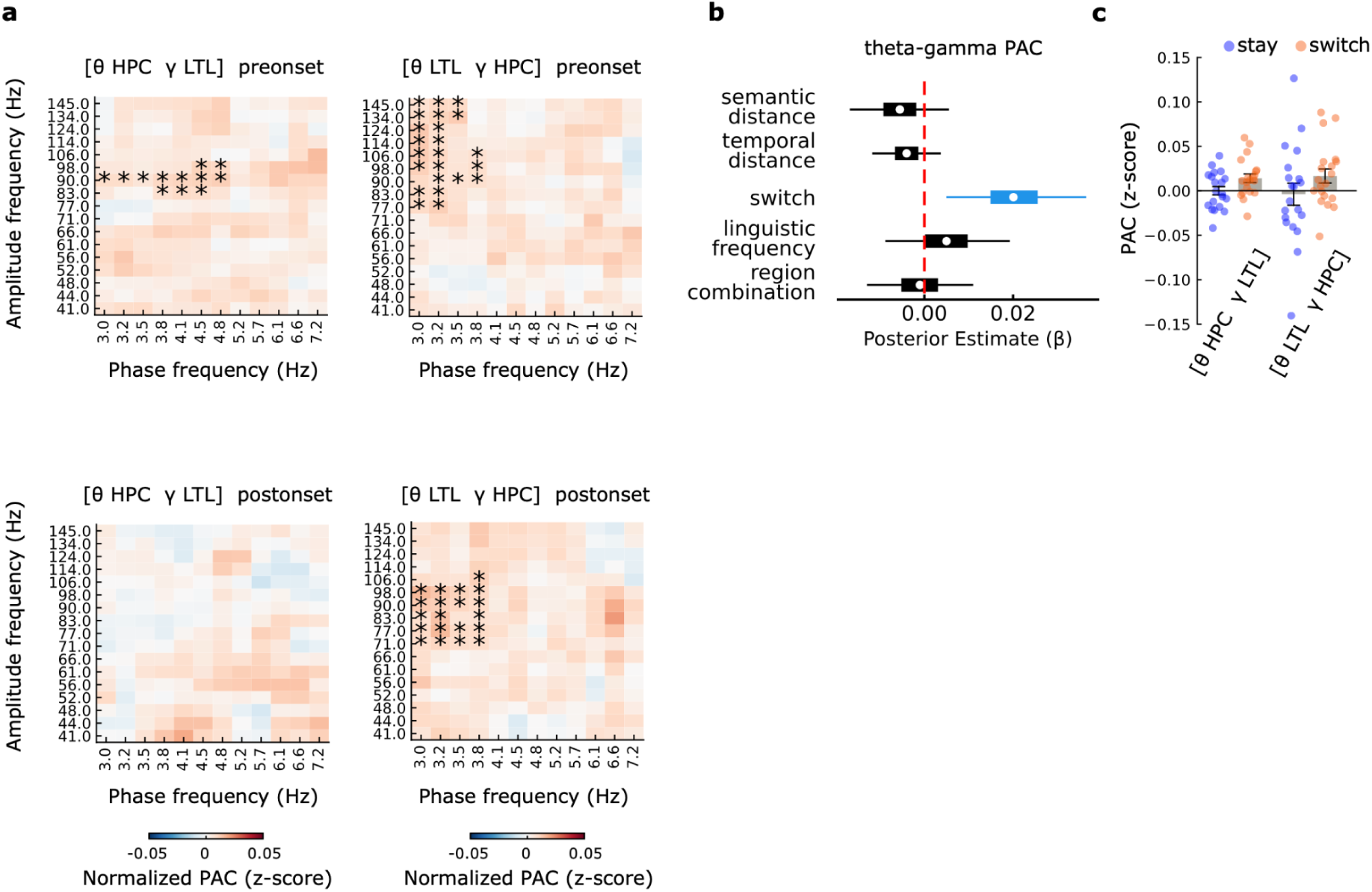
Cross-regional theta-gamma coupling between hippocampus and lateral temporal cortex supports semantic switching. **(a)** Theta-gamma PAC comodulograms during the pre-onset period (upper) shows significant cross-regional coupling between hippocampal theta phase and lateral temporal gamma amplitude (left), and vice versa (right). Asterisks mark clusters surviving cluster correction, indicating theta-gamma coordination between hippocampus (HPC) and lateral temporal lobe (LTL) during semantic retrieval. During post-onset period (bottom), this was observed only for phase of LTL theta and amplitude of hippocampal gamma. **(b)** Posterior estimates from a Bayesian linear mixed-effects model predicting cross-regional TG-PAC as a function of semantic (β = −0.006, 94% HDI = [−0.017, 0.006]) and temporal distance (β = −0.004, 94% HDI = [−0.012, 0.004]), lexical frequency (β = 0.005, 94% HDI = [−0.009, 0.019]), switch status (β = 0.02, 94% HDI = [0.005, 0.036]), and region combination (β = −0.001, 94% HDI = [−0.013, 0.011]). Semantic switching (blue) uniquely modulated cross-regional TG-PAC. **(c)** Visualization of TG-PAC strength as elevated for “switch” trials (orange) compared to “stay” trials (blue) across both phase-amplitude pairings. Dots represent individual participants. Error bars represent standard error of the mean.

Taken together, our results suggest that cluster transitions during internal mental search in conceptual spaces may involve coordinated dynamics across hippocampus and regions of the semantic network.

## Discussion

Internally directed search, whether for memories, words, or ideas, is a core feature of human cognition. Here, we investigated the role of hippocampal oscillations in guiding internal search. Using direct intracranial recordings during a verbal fluency task, we found that hippocampal theta systematically tracked semantic search: theta power increased with the conceptual distance between successive words that participants were spontaneously recovering from their knowledge. Crucially, participants exhibited a tendency to recover words that were similar in meaning close to each other and, occasionally, they switched to new groups or “clusters” of semantically similar words, similar to exploratory shifts between resource patches when animals forage in the external physical world. This tradeoff between exploration and exploitation was marked by stronger theta-gamma phase-amplitude coupling (TG-PAC) both within the hippocampus and across the hippocampus and the lateral temporal cortex, which hosts a key node of the semantic and language networks. These findings suggest that internal foraging for concepts in memory might rely on dynamic hippocampal-cortical coordination, and that this might be governed by oscillatory mechanisms that closely mirror those used during spatial exploration.

Our findings are in line with the idea that neural mechanisms supporting spatial navigation also support the structuring and the manipulation of abstract knowledge, aligning with proposals that the hippocampus facilitates cognitive mapping across diverse domains (Bellmund et al., 2018; Bottini & Doeller, 2020; Buzsáki & Moser, 2013). Prior work has demonstrated relational coding in the medial temporal lobe using fMRI across a variety of conceptual spaces, including visual features (Constantinescu et al., 2016; Theves et al., 2019, 2020; Viganò et al. 2023), odors (Bao et al., 2019), audiovisual and linguistic material (Viganò & Piazza 2020; Viganò et al. 2021), social hierarchies (Park et al., 2021), value (Nitsch et al., 2023), mathematical operations (Eperon et al., 2025), action plans (Barnaveli et al. 2025), and emotion (Qasim et al., 2023). These previous experiments, however, mostly relied on laboratory-constructed conceptual spaces or episodic recall of predefined items. In the present study, on the contrary, we used a verbal fluency task that offered a more naturalistic and ecologically valid window into cognitive maps, where people could spontaneously sample concepts from their knowledge about the world. This revealed that hippocampal theta power scale with conceptual distance during such spontaneous semantic search, and that hippocampal activity dynamically coordinates internally and externally (with lateral temporal regions) in supporting transitions between conceptual subspaces, via theta-gamma phase amplitude coupling. This extends recent fMRI findings (Lundin et al., 2023) by identifying temporally precise electrophysiological signatures of semantic exploration and connecting it to a network perspective. Moreover, we recently showed that verbal fluency tasks are accompanied by spontaneous eye movements that reflect several properties of the explored conceptual space (Viganò et al. 2024; Viganò, Cristoforetti et al. under review). An intriguing perspective, given the existing link between eye movements and attention (Rizzolatti et al., 1987; Corbetta et al., 1998; Smith et al., 2012; Awh et al., 2006), is that internal attentional shifts may be the way by which conceptual spaces are navigated. Indeed, the neurocognitive machinery implicated in cognitive maps, such as entorhinal grid-like coding, has been shown to activate both as a function of eye movements and covert attentional shifts (Killian et al. 2012; Nau et al. 2018; Wilming et al. 2018; Giari et al. 2023). Future experiments will be needed for properly understanding the link between attention, cognitive maps, and the oculomotor system.

Our results are substantially different from a recent intracranial study by Solomon et al. (2019), which linked hippocampal theta activity to semantic distances between words recalled from memorized lists. While both studies implicate hippocampal theta in relational processing during verbal retrieval, there are key differences. First, Solomon et al. focused on episodic memory, as they asked participants to recall previously encoded sequences of unrelated words, whereas our task targeted spontaneous semantic fluency, tapping into long-term semantic stores rather than temporally bound episodes. Second, while Solomon et al. observed stronger effects for low-dimensional semantic embeddings, we found that hippocampal theta more robustly tracked high-dimensional distances, potentially consistent with the idea that semantic memory may operate in a higher-dimensional representational space (Lund & Burgess 1996; Piantadosi et al., 2024). This raises the intriguing possibility that the hippocampus may construct lower-dimensional maps for episodic sequences, but navigate richer spaces during semantic search. Third, we observed robust high-frequency gamma activity and theta-gamma phase-amplitude coupling, as well as synchronization with lateral temporal lobe around cluster switches, features absent in Solomon et al., highlighting the temporal dynamics and interregional coordination that underlie exploratory semantic search. Together, these distinctions suggest that the hippocampus supports flexible retrieval strategies across memory systems, using domain-sensitive oscillatory mechanisms beyond episodic memory.

Still, the role of the hippocampus in semantic memory remains debated, but growing evidence points to its involvement in semantic retrieval (Duff et al., 2020). Prior studies using language tasks have shown increases in hippocampal theta (Piai et al., 2016) and gamma (Jafarpour et al., 2017) power when contextual cues facilitate semantic access. Building on this, we observed a striking temporal profile of hippocampal theta power during our verbal fluency task: an initial increase, a peak around 0.4 s before word onset, followed by a decline. This triphasic pattern might mirror (in reversed order from stimulus onset, given our task) the stages proposed in psycholinguistic models of speech production, such as Indefrey and Levelt’s (2004), which distinguish conceptual preparation, lexical access, and phonological encoding. We speculate that in our task, the early theta rise might reflect the conceptual search phase, the peak might mark lexical access upon finding a concept, and its decline might signal a transition to language-specific processes beyond the hippocampus. While this interpretation remains tentative, it opens avenues for linking oscillatory dynamics to functional stages of semantic foraging. Importantly, our findings also highlight dynamic coordination between the hippocampus and language-related cortical areas, particularly the lateral temporal lobe (LTL), a core node in the semantic network (Binder et al., 2009; Fedorenko et al., 2024). While the hippocampus may not store semantic content per se, it may act as a pointer system that accesses and coordinates representations distributed across cortical regions (Teyler & DiScenna, 1986; Teyler & Rudy, 2007; Simons et al., 2022). Supporting this, prior work has shown widespread hippocampal-cortical synchrony (Sirota et al., 2008; Gattas et al., 2023) and gamma coherence between hippocampus and LTL during spatial memory tasks (Pacheco Estefan et al., 2019). In particular, cross-frequency coupling, where theta phase aligns across distant regions and gamma power synchronizes locally, has been proposed as a mechanism for overcoming long-range transmission constraints (Hyafil et al., 2015; Buzsáki & Schomburg, 2015; Griffiths & Jensen, 2023). Our findings of increased theta-gamma coupling and hippocampal-LTL synchrony during semantic cluster switches are consistent with this framework, reinforcing the view that the hippocampus acts as a dynamic coordinator of conceptual search within a distributed cortical network. What cognitive phenomenon might trigger the switch to a new cluster remains unclear. In our behavioral analyses we isolated a potential crucial role played by linguistic frequency, as cluster switches are typically preceded by a significant drop of this parameter. However, we did not observe any significant modulation of hippocampal (or LTL) oscillatory power related to this aspect, stressing the necessity of further investigating it via more dedicated experiments or accessing other brain regions.

Finally, we have observed a significant decrease in power in alpha and beta frequency bands which, however, we couldn’t relate to any of the behavioral effects we isolated in participants’ performance. The temporal extent and time position of the alpha event, in particular, seems precisely aligned with the decrease in gamma and an increase in theta, suggesting a potential link between these changes. At the present moment, however, we can only speculate on whether these events might be related to more basic processes (e.g., motor articulation), or they might reflect unknown internal mechanisms for which further experiments will be needed.

Altogether, our results shed new light on the neural dynamics that support internal search through semantic memory. By demonstrating that hippocampal theta tracks the distance between concepts during spontaneous semantic search, and that both hippocampal and interregional theta-gamma coupling with LTL increases during exploratory transitions of semantic memory, we provide a better mechanistic understanding of how internal foraging might be supported by the brain. As such, this work suggests the hippocampus as a domain-general “search engine”, dynamically guiding access to meaning across our mental landscape.

## Acknowledgments

SV’s research is supported by the Max Planck Society. GG’s and RB’s research is supported by ERC-StG, NOAM 804422, and ERC-CoG, ATCOM, 101125658, both attributed to RB. RM’s research is supported by Hospital Niguarda ‘Ca Granda, Milan. CFD’s research is supported by the Max Planck Society and the Kavli Foundation. We thank the personnel at the “Claudio Munari” Center for Epilepsy and Parkinson Surgery, Niguarda Ca’ Granda Hospital, Milan, Italy, and the patients who agreed to participate in this experiment. We thank Mattia Silvestri and Francesco Romandini for help with data collection. Moreover, we thank our lab colleagues, in particular Nicholas Menghi, Qiaoli Huang, Marit Petzka, Irina Barnaveli, Alex Eperon, and Federica Sigismondi, as well as Tobias Staudigl and Virginie van Wassenhove, for insightful comments and suggestions.

## Authors’ contribution

SV and RB conceived the study, its research question and the experimental paradigm, with inputs from CFD and coordinating with RM for its implementation. SV and GG collected data, with support from RM. GG analyzed data, with input from SV and RB. All the authors discussed the results and their interpretation. SV and GG wrote the manuscript draft. All the authors contributed to the final version of the manuscript.

## Data and code availability statement

Due to their clinical nature, preprocessed and anonymized data will be shared upon request, in compliance with the internal policy of Niguarda Hospital.

Analysis code is available at https://github.com/giulianogiari/SemanticForaging

## Methods

### Participants

Twenty-one patients with drug-resistant epilepsy were recruited for the study. Patients underwent surgery and were implanted with intracranial stereotactic-electroencephalography depth electrodes (sEEG, Nihon-Koden) for seizure monitoring and localization of epileptic regions. All patients had at least one contact in the hippocampus. Additional information about the patients can be found in Table 1. The study was approved by the ethical committee of the University of Trento and of the Niguarda Hospital. One participant was excluded due to clinical assessment (extensive epileptic activity) by RM.

### Experimental design

Participants performed a categorical verbal fluency task for three different semantic categories (animals, professions, cities). The name of a category was presented on a computer screen for three seconds, followed by a 0.1 s long sound cue indicating the start of the task block. Participants had then two minutes to name as many concepts as possible for the given category. They were instructed to speak calmly, without rushing to avoid creating overlap between words. During these two minutes a green fixation cross was presented on a black screen, after which the fixation cross turned white and another sound cue (0.1 s) instructed participants to rest for one minute. In a first run, this alternation of verbal fluency task and rest was repeated three times, one for each category with the order of the categories randomized across participants (Figure 1b). In a second run instead, after the resting period, participants were presented again with the same category and were instructed to retrieve even more concepts than in previous blocks, with the possibility to repeat concepts that were already being said. This manipulation was introduced to address other scientific questions that are not part of the current investigation.

Before the main experimental procedure with the three aforementioned categories, participants familiarized themselves with the verbal fluency task with the category “objects”, being asked to mention for two minutes all the objects they could think of. This block was not recorded and was thus not used in the analyses.

Visual and auditory presentation, recording of participants’ voices and control of experimental timings were performed using MATLAB and Psychtoolbox (Brainard, 1997). Lights were switched off during the experiment so that the room was dimly lit to minimize the influence of external visual information.

One participant (Participant ID 1) did not do the task for the “cities” category, thus for this participant only the data of the other two categories were used.

### Experimental events definition

Audio recordings of the participants were segmented using Audacity (https://www.audacityteam.org/). The generated time stamps were visually inspected and labelled with the pronounced word by SV and GG. Time stamps of each word were realigned to the trigger sent during the sEEG recording at the beginning of the corresponding foraging block. Epochs were defined as the time window starting 1.5 s before word onset and ending 1.5 s after word onset. Within these 1.5 s windows only 1 s was of interest, 0.5 s were included to account for edge artefacts caused by time-frequency analysis (Cohen, 2014; see later). Additional epochs of interest during the foraging block were defined based on silence periods between words. We considered only silence periods at least 5 s long. These time periods were further segmented in 2 s long time windows, to match the duration of the word epochs including 0.5 s to account for edge artefacts. Similarly, the resting periods were segmented in 2 s long time windows.

We excluded from all analyses the epochs that had an overlap of less than 1.5 s to avoid contamination between the neural activity in the period of interest (pre-word onset) and speech related activity of the previous word. This led to the exclusion of ∼28% trials. See Table 1 for the total number of words per participant included in the analyses. Note that distance metrics used for the analyses (see later) were calculated based on the word sequence as said by the participant before word exclusion, thus reflecting the actual navigation process of the participant.

### Behavioral descriptive statistics

We evaluated participants’ behavior by calculating the amount of words produced in each verbal fluency task block as well as their inter-word time interval. We used a series of one-way analysis of variance (ANOVA) to compare the amount of words recalled between semantic categories and between repetitions of the task. Similarly, we compared the inter-word time interval across categories and across repetitions. These first analyses provided a descriptive measure of participants’ exploration behavior.

### Evaluating semantic distances as predictors of behavior

To investigate which distance metric better accounted for participants’ timing behavior, we obtained, for each word, several language-related variables and used them to compute a distance metric between subsequent words as spoken by the participants.

Specifically, we used: i) word embeddings from the Italian FastText (Bojanowski et al., 2016), both in their original dimensionality (300) and, following previous studies (Solomon et al., 2019), reduced to a lower dimensionality (1, 2, 3 dimensions) using principal component analysis as implemented in scikit-learn (Pedregosa et al. 2011). As distance metric we used cosine for the high-dimensional vector and Euclidean for the low dimensional vectors; ii) word frequency, measured as Zipf value obtained from the wordfreq package (https://zenodo.org/records/7199437). Distance was defined as the difference in frequency scores; word length, i.e., number of letters, and the distance is their difference; path distance obtained from the wordnet corpus (Miller, 1995) as implemented in the nltk package (Bird 2009).

Each distance measure was then correlated with the time interval between subsequent words. The rationale was that, assuming constant speed, the time interval is the best indicator of traveled distance. The linguistic variable that correlates better with inter word interval can thus be interpreted as better reflecting the underlying conceptual space. This correlation was repeated for each participant, category and task block and finally averaged to obtain one correlation value per participant. Correlation values were fisher transformed and tested against zero using a two-tailed, one-sample t-test. We further tested whether the highest correlation was higher than the others using a two-tailed, paired sample t-test. The correlation between the high-dimensional semantic vectors and ITI was repeated after removing the “switch” trials (see later) and the resulting fisher transformed correlation values tested against zero using a two-sided, one-sample t-test.

Using the distance metric that is best correlated with timing intervals between words (i.e., high-dimensional word embeddings), we evaluated whether it is capturing meaningful aspects of participants’ sequence of named concepts. To this end, the word sequence of each task block of each participant was randomly shuffled 1000 times and the distance between subsequent words was recomputed and averaged to obtain 1000 distance scores per participant. We then normalized the observed average distance between sequent words by the null distribution of 1000 average distance scores. The obtained z-scores were tested at the group level against zero using a two-tailed, one-sample t-test.

### Behavioral signatures of foraging in conceptual spaces

We evaluated whether naming concepts in the verbal fluency task mimicked strategies that animals use when foraging in physical space. Specifically, we tested the presence of local clusters of concepts that are visited in sequence (exploitation) followed by “switches” between clusters (exploration). We used the “similarity drop” model (Hills et al., 2012; Lundin et al. 2023) as implemented in the “forager” package (Kumar et al., 2024) to define “switch” trials as the points where semantic distance between subsequent words is higher than its neighbors. Words in between switch trials were considered to belong to the same “cluster”. In order to assess whether the so-defined clusters are meaningful, we tested whether distances “within” clusters were smaller as compared to distances “between” clusters. This comparison was by design unbalanced due to the low amount of switch trials, and potentially circular in that the distance we evaluated was used to define the clusters. To overcome these limitations we created a null distribution of clusters and their “within” and “between” distances by randomly shuffling 1000 times the order of words within each task block and redefining the switch trials at each iteration. The observed “within” and “between” distances were then normalized based on the null distribution of the respective distances. The obtained z-scores were then tested at the group level using a two-tailed, paired-samples t-test.

After having observed that participants have a tendency to exploit local patches before moving on to explore other patches in the environment, we wanted to evaluate how participants choose the next patch to explore. Based on the literature (e.g., Hills et al., 2012), we evaluated whether linguistic frequency changes as a function of its position with respect to the cluster switch. To this end we extracted the linguistic frequency (Zipf value) of each word around the cluster switch. Then, for each participant, we averaged the Zipf score of the different words belonging to the same position with respect to cluster switch. We next tested at the group level whether the frequency of subsequent orders (e.g., −2 vs −1 relative to cluster switch) differed statistically using a two-tailed, paired sample t-test.

### Intracranial stereotactic EEG recording and preprocessing

sEEG recordings were performed at the Niguarda Hospital (Milan, Italy) using a Nihon-Koden system with a sampling rate of 1000 Hz. Data were recorded with an online reference chosen for clinical reasons. Raw data were visually inspected by expert epileptologist (R.M.), who indicated contacts and time windows contaminated by potential epileptic discharges that were subsequently marked for exclusion. Additionally, we evaluated the amount of data excluded against automated approaches (Gelinas et al., 2016) demonstrating that a comparable percentage of data is discarded (Supplementary Figure 9). Raw sEEG files were imported in Fieldtrip (Oostenveld et al., 2010) and converted to BIDS format (Gorgolewski et al., 2016). BIDS-formatted data were then loaded in python and data analysis was further carried out using MNE-python (Gramfort et al., 2013) as well as common scientific python packages. Time series data were low-pass filtered at 150 Hz to avoid aliasing artefacts (Cohen, 2014) and notch filtered at 50±1, 100±1 and 150±1 Hz (line-noise and harmonics). Data were re-referenced offline using the “bipolar” reference scheme, which subtracts the activity of neighboring contacts from medial to lateral along the sEEG electrode, effectively increasing the spatial specificity of the recordings (Mercier et al., 2022). Neighboring contacts that were localized in different brain areas based on the Desikan-Killiany atlas (Desikan et al., 2006) were not included in further analyses to avoid misinterpreting the spatial origin of the effects. After epoching based on events onset (see paragraph “Experimental events definition”), data were downsampled to 500 Hz to ease computational cost of the following analyses.

### Electrode localization

Electrode localization was performed based on coregistered post-implant CT and pre-implant MRI. Anatomical electrode localization was based on the Desikan-Killiany atlas (Desikan et al., 2006) and confirmed through visual inspection of the electrical signal by an expert epileptologist (R.M.). The following analyses were performed on contacts localized in the hippocampus. Furthermore, we defined two control regions, one in the lateral temporal lobe, which is known for its role in language processing and semantic memory (Binder et al., 2009; Fodorenko et al. 2024) and one in the precentral gyrus which, on the contrary, should not be involved in any of these processes. Lateral temporal contacts were defined as the grey-matter contacts on the same shaft of the hippocampal contacts and were present in all the participants, while precentral contacts were missing in 9 patients.

### Time-frequency analysis

Single-trial time series were convolved with Morlet wavelets to obtain time-resolved, frequency-specific power estimates. Fifty Morlet wavelets were constructed with logarithmic spacing between 3 and 145 Hz. Line noise and its harmonics were at least 1 Hz away from the closest wavelets. Wavelet cycles were frequency dependent and ranged between 3 (lowest frequency) and 5 (highest frequency). To avoid edge artifacts due to time-frequency decomposition (Cohen, 2014), the 0.5 s at the beginning and at the end of the trial were excluded from further analyses after wavelet decomposition. Following (Tallon-Baudry & Bertrand, 1999; Iemi et al., 2017; Ronconi et al., 2017) we evaluated the temporal resolution of the selected time-frequency parameter: temporal resolution for the slowest frequency we considered (3 Hz) was of ∼0.160 s, while for the fastest (145 Hz) it was 0.05 s. Interpretation of effects within these latencies should thus be taken with caution as it can result from temporal smearing due to time-frequency decomposition. This is particularly relevant at the edges of the time-frequency spectrum, where effects can actually spread from neighbouring events, such as speaking. Power values were then log transformed given their chi-squared distribution (Percival, 1993; Manning et al., 2009).

Before subsequent analysis we used irregular-resampling auto-spectral analysis (IRASA, Wen & Liu, 2016) to obtain an estimate of the aperiodic component of the power spectrum. The IRASA algorithm proceeds by resampling (both upsample and downsample) the time-series of single trials with multiple time-warping factors (we used 13 linearly spaced factors from 1.1 to 1.75 in steps of 0.05) and recompute the time-frequency spectrum (same parameters as above). For each resampling factor we computed the geometric mean of the upsampled and downsampled time-averaged, frequency representation. The median across all resampling factors is an estimate of the aperiodic component, i.e., the non-oscillatory part of the signal. This was log-transformed and subtracted, at each time point, from the mixed time-frequency representation obtained using Morlet wavelets.

This procedure was repeated separately for the time window prior to word onset (preonset), the time window after starting pronouncing the word (postonset), the time window after word offset (postoffset), the periods of silence between words during the foraging task (silence) and the resting periods between foraging blocks (rest)

### Statistical analysis of time-frequency

Power in the time window before word onset was baseline-corrected with the average power in the time window after word onset. We then used a two-sided, one-sample t-test against zero at each time-frequency point. Multiple comparison correction was performed using false discovery rate (FDR, Genovese et al., 2002). This analysis was repeated for both the hippocampus and the control regions.

As control analyses we evaluated power differences between baselines, categories and repetitions of the same category. We selected frequencies in the significant time-frequency ranges identified in the main contrast and averaged their power across frequencies and time points. We thus obtained estimates of power for each frequency band: theta: 3-7.2 Hz, −1 to −.2 s; gamma 41-145 Hz, −0.968 to −0.444 s; alpha: 9.8-11.5 Hz, −0.668 to −0.572 s; beta: 15.8–34.8 Hz, −0.398 to −0.124 s. We then ran three one-way ANOVAs: one with factor baseline (four levels: postonset, postoffset, resting, silence); one with factor category (three levels: cities, animals, professions); one with factor repetition (three levels: 1,2,3). For the silence and rest periods, we randomly selected an amount of segments that matched the amount of words being said by the individual participant (after trial exclusion based on temporal overlap, see previous paragraph).

### Identifying peaks in frequency-band power

To visualize their time course, we averaged the power in the significant frequency bands identified in the time-frequency analysis (theta: 3-7.2 Hz; gamma 41-145 Hz; alpha: 9.8-11.5 Hz; beta: 15.8–34.8 Hz). We identified the peak in the respective frequency band by considering the highest (for theta and gamma) or lowest (for alpha and beta) time point in the time series. We then evaluated the stability of the peak across participants using a jackknife procedure often employed to evaluate the peak-latency of event-related potentials (Miller et al., 1998). The jackknife procedure involves removing one participant from the group and repeating the peak identification procedure. The procedure was repeated until all participants were left out once. This allowed us to obtain a distribution of peak estimates from which we computed the 95% confidence interval.

### Theta-Gamma phase amplitude coupling (TG-PAC)

We estimated the consistency with which the amplitude of gamma increased at specific phases of theta, i.e., theta-gamma phase-amplitude coupling (TG-PAC). We defined theta as 12, log-spaced frequencies between 3 and 7.2 Hz, while gamma was defined as 16 log-spaced frequencies between 41 and 145 Hz. These were the same frequencies used for the time-frequency analysis. For each frequency, the filter range was defined as ±(center frequency ÷ 4) for theta phase and ±(center frequency ÷ 8) for gamma amplitude, following Bahramisharif et al. (2013). The data were then band-pass filtered in these ranges and the Hilbert transform was applied to obtain the instantaneous phase (theta) and amplitude (gamma). To avoid edge artifacts from the filtering process, we removed 0.5 s from the start and end of each segment. The analysis focused on the preonset and postonset time windows.

TG-PAC strength was quantified using the modulation index (MI; Tort et al., 2010), resulting in a TG-PAC value for each trial and contact. To estimate the reliability of observed TG-PAC values and control for spurious coupling, we generated a null distribution by randomly shuffling phase and amplitude pairs across trials 200 times (Tort et al., 2010). The observed TG-PAC values were then z-scored using the mean and standard deviation of this surrogate distribution.

This analysis was repeated considering phase and amplitude estimated on the same contact, thus being informative about within-region TG-PAC, as well as estimating theta phase on a contact in a given region (e.g., hippocampus) and the amplitude of a contact in another region (e.g., LTL). This latter procedure was repeated for all possible combinations of contacts across the two regions and was informative about the strength of between-region TG-PAC.

All the TG-PAC analyses were carried out using the tensorpac toolbox (Combrisson et al., 2020).

### Statistical analysis of TG-PAC strength

To assess which phase-amplitude frequency pairs showed significant TG-PAC at the group level, we averaged contacts for each participant, and tested the null-hypothesis of no-PAC using a cluster-permutation test to correct for multiple comparisons across the phase frequency x amplitude frequency comodulogram. The cluster-permutation test first performed a one-sample t-test at each frequency x amplitude pair. If contiguous pairs survived the uncorrected threshold of p < 0.05, they formed a cluster. The observed cluster statistic was the sum of t-scores of the individual, adjacent pairs. This procedure was repeated 10000 times, each time shuffling the sign of the z-scored TG-PAC value. At each iteration we retained the highest cluster statistic to create a permutation distribution to which the observed cluster statistic was compared to eventually derive a p-value corrected for multiple comparisons. This analysis was repeated to investigate the presence of significant TG-PAC within region and across regions.

### Control analyses on within-region TG-PAC

We evaluated whether the strength of the observed TG-PAC could be influenced by properties of the interacting frequency bands (Cole & Voytek, 2017; Aru et al., 2015). To this end we performed two control analyses: 1) we correlated the strength of the TG-PAC with theta and gamma power, separately, to investigate whether the observed increase in TG-PAC can be explained by a better estimation of frequency band power; 2) we computed two properties of theta waveform shape, the rise-decay asymmetry and the peak-through asymmetry using the bycycle toolbox (<u>Cole & Voytek, 2019</u>). If theta was a perfect sinusoid, its value in these metrics would be 0.5, we thus compared the observed values to 0.5 using a one-sample t-test as well as between the preonset and postonset time windows using a paired sample t-test.

### Linear-mixed models

Linear-mixed models are powerful tools to assess the contribution of an independent variable on the dependent variable when multiple correlated measurements are performed (e.g., from the same participant performing multiple trials and having multiple electrodes, Laird & Ware, 1982). We used linear-mixed models to investigate whether hippocampal activity was related to the relevant aspects of navigation behavior. Specifically, we wanted to evaluate whether changes in hippocampal activity are functionally related to (i) semantic distance between concepts, (ii) retrieval latency, (iii) balance between local exploitation and global exploration via semantic switches, and (iv) word frequency, all of which play a role in participants’ semantic foraging behavior.

We thus selected these predictors as previously defined (see above). The distance assigned as predictor to a given trial corresponds to the distance between the word pronounced in the same trial t and the previous one t-1. For example, if in trial t the participant pronounced the word “dog”, the predictor for trial t would be the distance between the word vector for “dog” and the word pronounced in trial t-1. The first word of each foraging block thus has no meaningful distance and was excluded from the analyses. Continuous predictors (semantic distance, temporal distance, lexical frequency) were z-scored given their different scales, allowing a more direct comparison of the resulting parameter estimates. Following Stangl et al., (2020), we proceeded on fitting separately a full model and a series of models with a single predictor (plus its random slope and intercept), allowing us to evaluate the impact of correlated predictors on the estimates. The full model had the following formula:

*data ∼ semantic distance + temporal distance + switch + linguistic frequency +*

*(1 + semantic distance + temporal distance + switch + linguistic frequency | subject)*

The same model was fit to different data. Data in the formula can be one of:

- power, the trial-level frequency-band specific power, selected from the TFR as being either theta (3-7.2 Hz, −1 to −.2 s), gamma (41-145 Hz, −0.968 to −0.444 s), alpha (9.8-11.5 Hz, −0.668 to −0.572 s), beta (15.8–34.8 Hz, −0.398 to −0.124 s). As a measure of power we used the difference between the preonset time window and the average of the baseline period (post-onset), after correction for the aperiodic component by IRASA;
- TG-PAC, the trial-level normalized TG-PAC estimate. Given our hypothesis on the increase of TG-PAC being related to cluster-switch behavior, this analysis focused on trials around the switch point, that is the trial labeled as switch and its previous trial. This ensures to have an equal amount of trials for both levels of the switch predictor.

The reduced model included only one predictor at the time and its corresponding random slope. Using semantic distance as an example, the model had the following formula:

*data ∼ semantic distance + (1 + semantic distance | subject)*

For theta power, we repeated the model fitting procedure after exclusion of the switch trials. The model(s) thus had the same formula(s) as above except for the switch predictor.

For hippocampal TG-PAC, we also fitted a model including the time window as a categorical predictor. The model had the following formula:

pac ∼ switch * time window + (1 + switch * time window | subject)

where the time window is a categorical predictor with two levels, preonset and postonset. For cross-regional HPC-LTL TG-PAC, we fitted a model including the region combination as a categorical predictor. The formula(s) were the same as above plus the addition of a categorical predictor “region combination” with two levels (theta HPC, gamma LTL; theta LTL, gamma HPC).

The model fitting procedure was carried out using both a bayesian and a frequentist approach.

The analyses using Bayesian models were carried out using the Bambi toolbox (Capretto et al., 2022). For all the predictors we used weakly informative priors with mean 0 and sigma 1. The model was fitted using four Markov Chain Monte Carlo No-U-Turn sampler (NUTS), with each chain having 1000 burn-in steps followed by 4000 draws. All parameters of the main effects converged across the four chains (r_hat < 1.1) and had more than 1000 valid samples. We considered a fixed effect to be significant if the 94% of the high-density interval (HDI) of the posterior did not include zero, following standard settings in the arviz toolbox (Kumar et al., 2019). The analyses using frequentist models was carried out using pymer4 (Jolly, 2018). Models were fitted using the optimx package and the bobyqa solver. Significance was evaluated using an ANOVA. Before model fitting, data underwent rank-based inverse normal transformation to ensure normality of residuals. We also evaluated whether the correlation in the predictors (see Figure 1d) can be problematic for the interpretation of the results. To this end we computed the variance inflation factor (VIF, Montgomery et al., 2012) of the fixed effects using statsmodels (Seabold and Perktold, 2010). The VIF measures the degree to which the variance explained by one predictor is inflated due to multicollinearity with other predictors in the model. A value higher than 5-10 warrants caution in the interpretation of the models estimate. VIF for all predictors was slightly above 1 (semantic distance: 1.274; temporal distance: 1.048; switch: 1.198; linguistic frequency: 1.023) indicating that multicollinearity was not affecting the parameter estimates of the model.

## Supplementary Materials

**Supplementary Figure 1.**
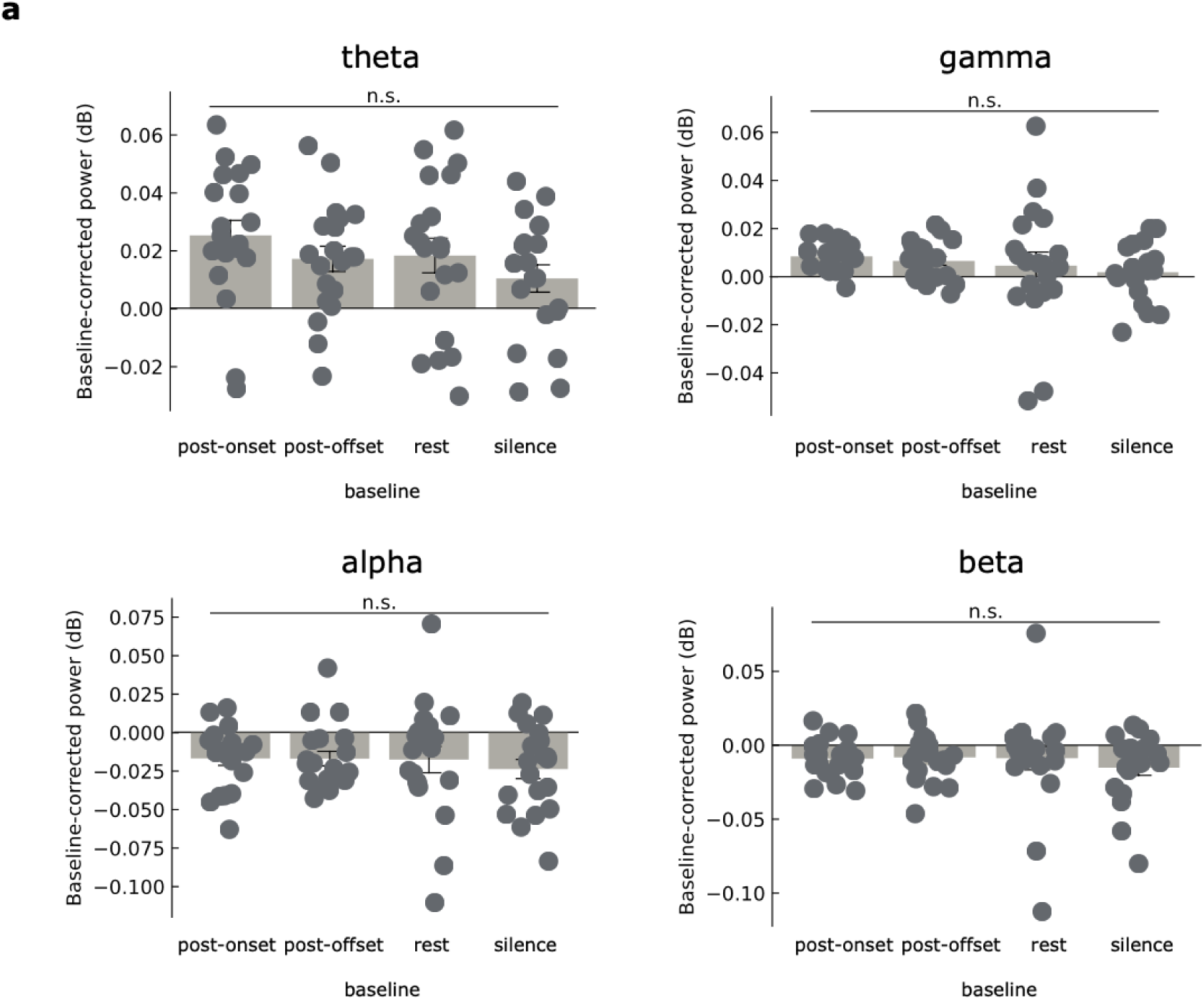
(a) We didn’t detect any statistically significant difference in hippocampal theta and gamma power increase, as well as alpha and beta decrease, as a function of different baselines. The different baselines were defined as follows: Post onset: 1 s after word onset; Post offset: 1 s after word offset; Rest: randomly sampled 1-s windows taken from “rest” blocks (see Methods); Silence: randomly sampled 1-s windows taken from periods of prolonged silence (at least 5 sec) between subsequent words (see Methods).

**Supplementary Figure 2.**
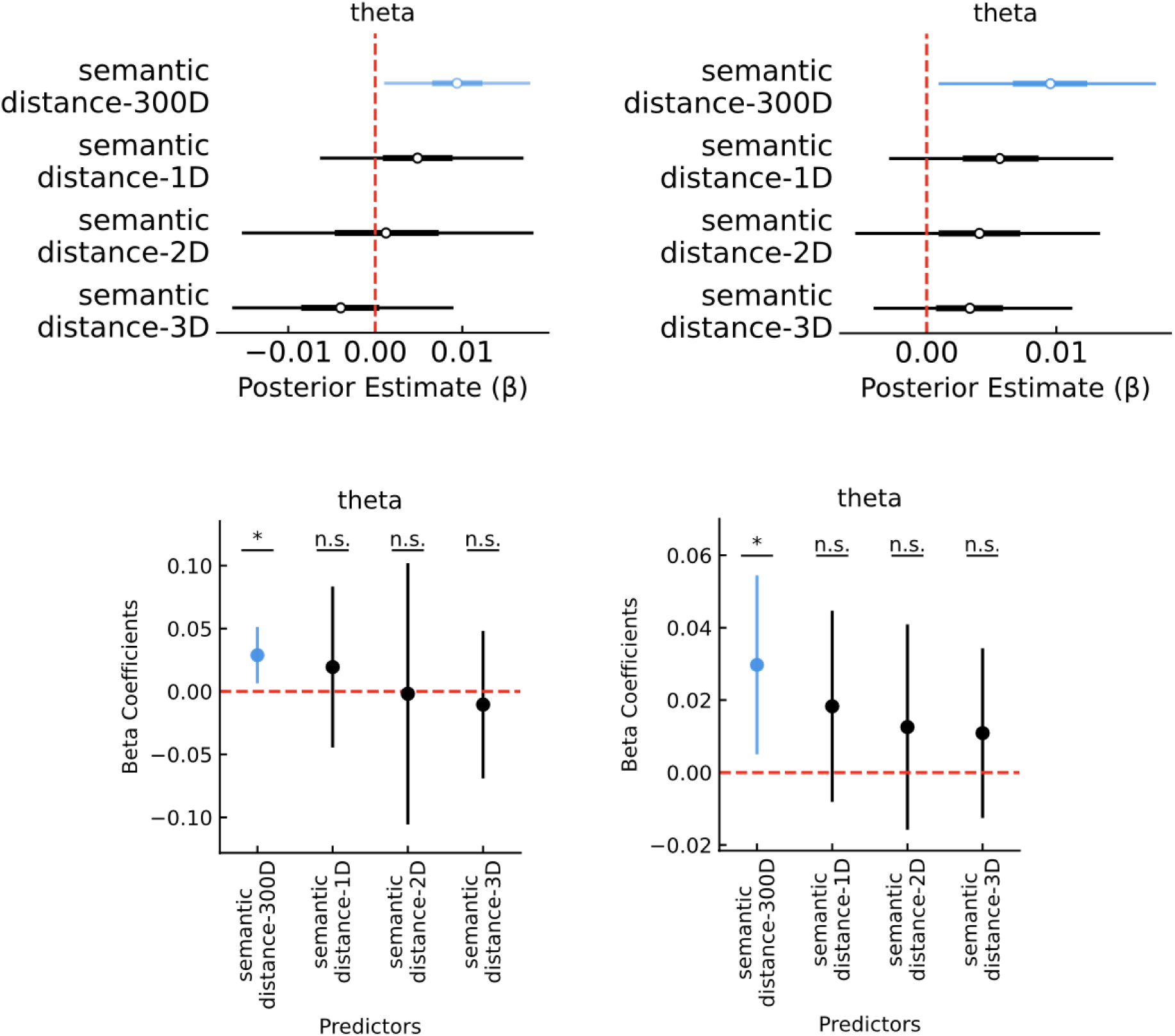
Linear mixed-effect models were conducted to verify whether semantic distances (Euclidean) in low-dimensional projections (via PCA) of fastText 300D space were still – or better – predicting changes in hippocampal theta power at a single trial level. We observed only a significant modulation of theta power by 300D semantic distance. We didn’t observe any statistically significant modulation as a function of 1D, 2D, or 3D distances. This effect was replicated using a variety of approaches> top left: Bayesian full LMM: 300D (β = 0.009, 94% HDI = [0.001, 0.018]), 3D (β =-0.004, 94% HDI = [−0.016, 0.009]), 2D (β = 0.001, 94% HDI =[−0.015, 0.018], 1D (β = 0.005, 94% HDI = [−0.006, 0.017]); top right: Bayesian individual LMM: 300D (β = 0.01, 94% HDI = [0.001, 0.018]), 3D (β = 0.003, 94% HDI = [−0.004, 0.011]), 2D (β = 0.004, 94% HDI = [−0.005, 0.013], 1D (β = 0.006, 94% HDI = [−0.003, 0.014]); bottom left: classical/frequentist full LMM: 300D (β = 0.028, CI= [0.006, 0.051], p=0.023), 3D (β =-0.01, CI= [−0.069, 0.048], p=0.732), 2D (β =-0.001, CI= [−0.1, 0.1], p=0.973), 1D (β =0.019, CI= [−0.044, 0.083], p=0.565); bottom right: classical/frequentist individual LMM): 300D (β = 0.029, CI= [0.005, 0.054], p=0.028), 3D (β = 0.01, CI= [−0.012, 0.034], p=0.374), 2D (β =0.012, CI= [−0.015, 0.04], p=0.397), 1D (β =0.018, CI= [−0.008, 0.044], p=0.19).

**Supplementary Figure 3.**
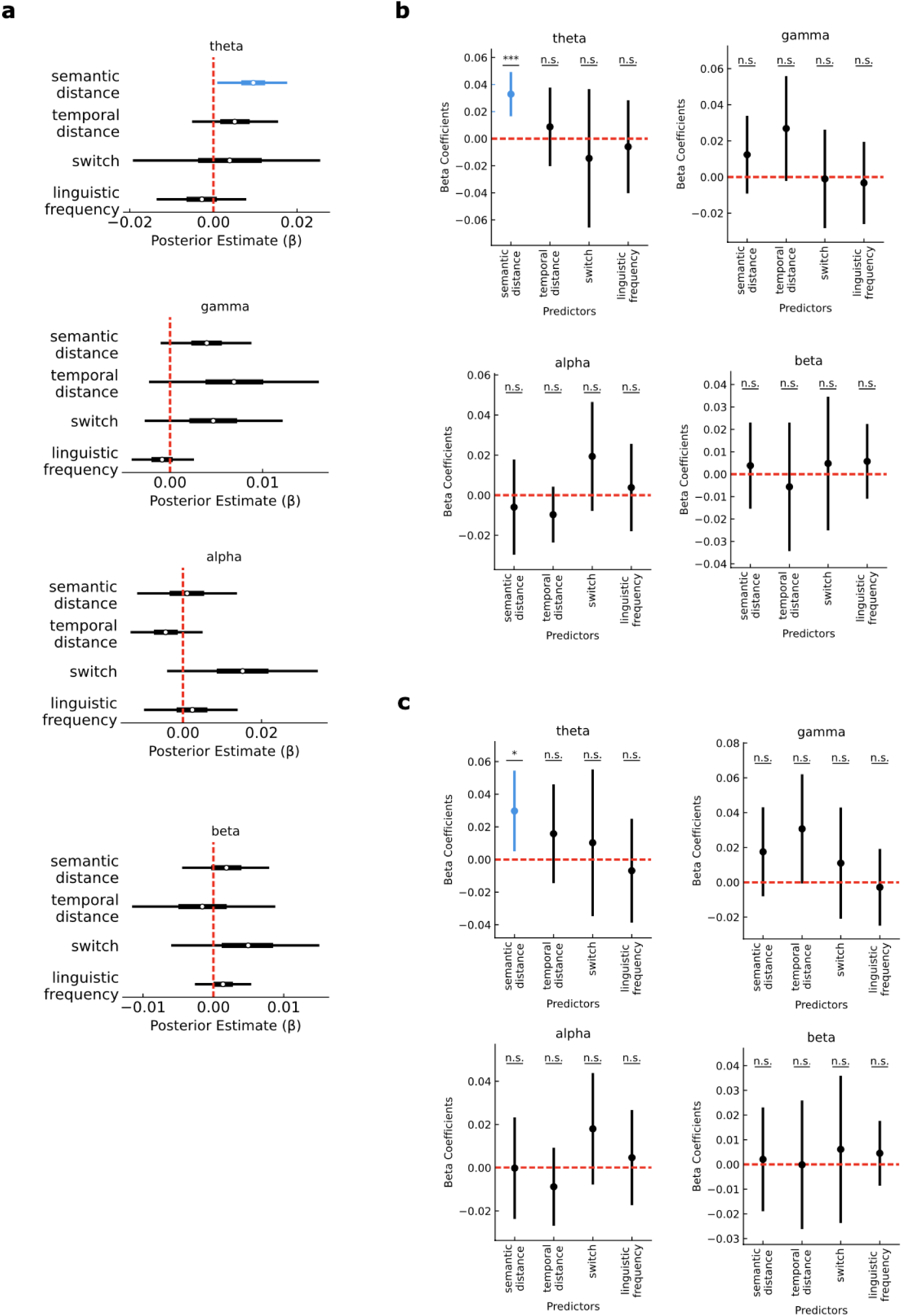
The effects reported in Figure 3 of main text (Bayesian full LMM) were robust across different modelling approaches. For all of them, we detected significant modulation of hippocampal theta power as a function of semantic distance at a single trial level. a) Bayesian individual LMM: theta: semantic distance (β = 0.01, 94% HDI = [0.001, 0.018]), temporal distance (β = 0.005, 94% HDI = [−0.005, 0.015]), switch (β = 0.004, 94% HDI = [−0.019, 0.026]), linguistic frequency (β = −0.003, 94% HDI = [−0.014, 0.008]); gamma: semantic distance (β = 0.004, 94% HDI = [−0.001, 0.009]), temporal distance (β = 0.007, 94% HDI = [−0.002, 0.016]), switch (β = 0.005, 94% HDI = [−0.003, 0.012]), linguistic frequency (β = −0.001, 94% HDI = [−0.004, 0.003]); alpha: semantic distance (β = 0.001, 94% HDI = [−0.012, 0.014]), temporal distance (β = −0.004, 94% HDI = [−0.013, 0.005]), switch (β = 0.015, 94% HDI = [−0.004, 0.034]), linguistic frequency (β = 0.002, 94% HDI = [−0.01, 0.014]); beta: semantic distance (β = 0.002, 94% HDI = [−0.004, 0.008]), temporal distance (β = −0.002, 94% HDI = [−0.012, 0.009]), switch (β = 0.005, 94% HDI = [−0.006, 0.015]), linguistic frequency (β = 0.001, 94% HDI = [−0.003, 0.005]); (b) classical/frequentist full LMM: theta: semantic distance (β = 0.032, CI= [0.016, 0.049], p< 0.001), temporal distance (β = 0.008, CI= [−0.020, 0.0377], p=0.565), switch (β = −0.014, CI= [−0.065, 0.036], p=0.585), linguistic frequency (β = −0.005, CI= [−0.040, 0.023], p=0.783); gamma: semantic distance (β = 0.012, CI= [−0.009, 0.033], p=0.280), temporal distance (β = 0.026, CI= [−0.002, 0.055], p=0.095), switch (β = −0.001, CI= [−0.028, 0.026], p=0.940), linguistic frequency (β = −0.003, CI= [−0.026, 0.019], p=0.781); alpha: semantic distance (β = −0.005, CI= [−0.029, 0.017], p=0.629), temporal distance (β = −0.009, CI= [−0.023, 0.004], p=0.185), switch (β = 0.019, CI= [−0.007, 0.046], p=0.176), linguistic frequency (β = 0.003, CI= [−0.017, 0.025], p=0.736); beta: semantic distance (β = 0.003, CI= [−0.015, 0.023], p=0.7), temporal distance (β =-0.005, CI= [−0.034, 0.023], p=0.707), switch (β = 0.004, CI= [−0.025, 0.034], p=0.757), linguistic frequency (β = 0.005, CI= [−0.010, 0.022], p=0.517). (c) classical/frequentist individual LMM: theta: semantic distance (β =0.029, CI= [0.005, 0.054], p=0.028), temporal distance (β =0.015, CI= [−0.014, 0.046], p=0.323), switch (β =0.010, CI= [−0.034, 0.055], p=0.660), linguistic frequency (β =-0.006, CI= [−0.038, 0.024], p=0.677); gamma: semantic distance (β = 0.017, CI= [−0.008, 0.043], p=0.195), temporal distance (β =0.030, CI= [−0.0006, 0.062], p=0.193), switch (β = 0.010, CI= [−0.020, 0.042], p=0.507), linguistic frequency (β = −0.002, CI= [−0.024, 0.0191], p=0.8); alpha: semantic distance (β = −0.0001, CI= [−0.023, 0.023], p=0.988), temporal distance (β = −0.008, CI= [−0.026, 0.009], p=0.374), switch (β = 0.018, CI= [−0.007, 0.043], p=0.190), linguistic frequency (β = 0.004, CI= [−0.017, 0.026], p=0.684); beta: semantic distance (β = 0.002, CI= [−0.018, 0.023], p=0.849), temporal distance (β = −0.001, CI= [−0.026, 0.025], p=0.992), switch (β =0.006, CI= [−0.023, 0.035], p=0.693), linguistic frequency (β = 0.004, CI= [−0.008, 0.017], p=0.501).

**Supplementary Figure 4.**
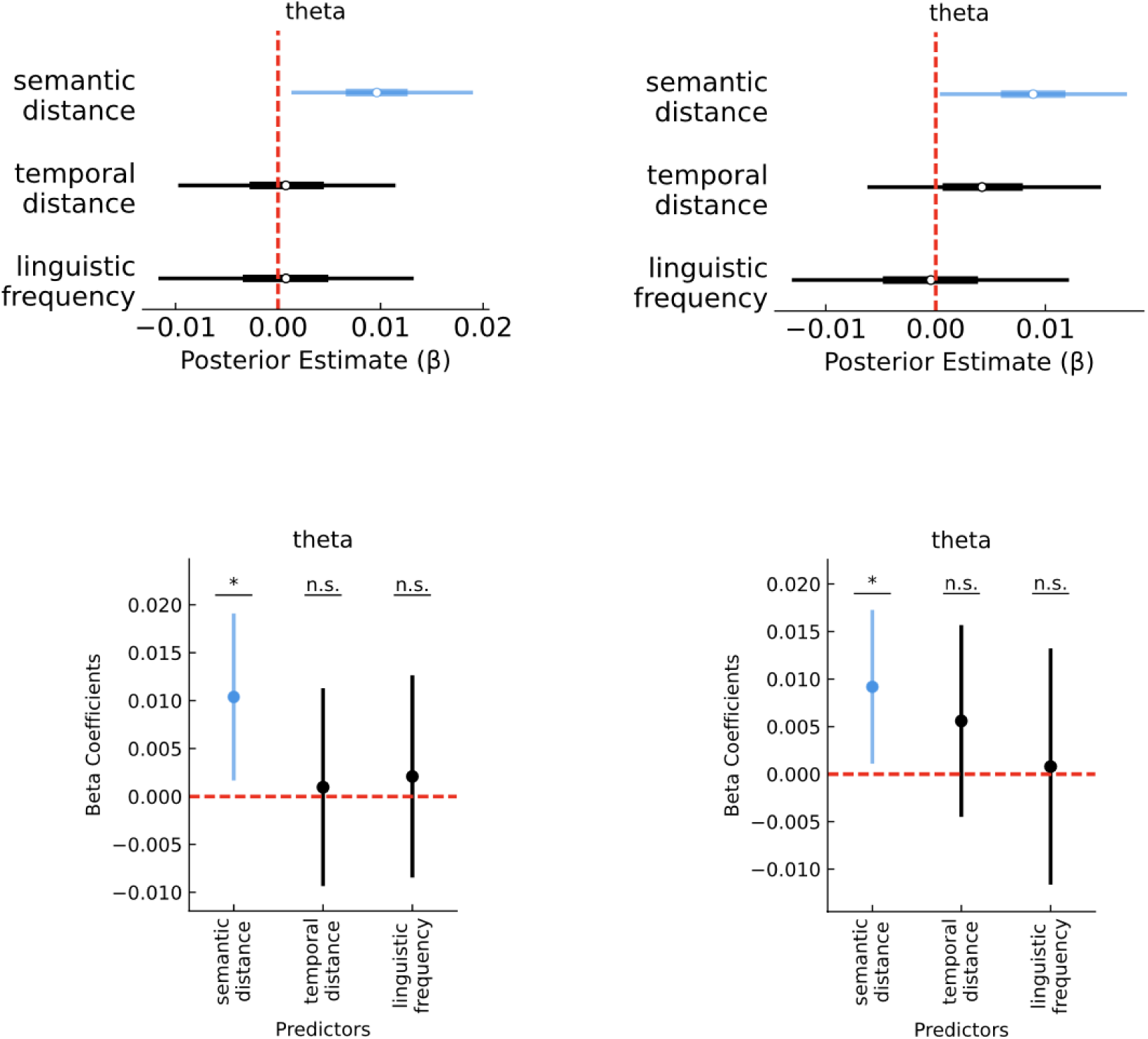
The effect reported in Figure 3a of main text (Bayesian full LMM) was robust across different modelling approaches also when restricted to “stay” trials, ignoring “switch” trials.(Top left: Bayesian full LMM: semantic distance (β = 0.01, 94% HDI = [0.001, 0.019]), temporal distance (β = 0.001, 94% HDI = [−0.01, 0.011]), linguistic frequency (β = 0.001, 94% HDI = [−0.012, 0.013]); top right: Bayesian individual LMM: semantic distance (β = 0.009, 94% HDI = [0, 0.017]), temporal distance (β = 0.004, 94% HDI = [−0.006, 0.015]), linguistic frequency (β = 0, 94% HDI = [−0.013, 0.012]); bottom left: classical/frequentist full LMM: semantic distance (β = 0.010, CI= [0.001, 0.019], p=0.029), temporal distance (β = 0.009, CI= [−0.009, 0.011], p=0.855), linguistic frequency (β = 0.002, CI= [−0.008, 0.0126], p=0.703); bottom right: classical/frequentist individual LMM: semantic distance (β = 0.009, CI= [0.001, 0.017], p=0.037), temporal distance (β = 0.005, CI= [−0.004, 0.015], p=0.293), linguistic frequency (β = 0.0007, CI= [−0.011, 0.013], p=0.901).

**Supplementary Figure 5.**
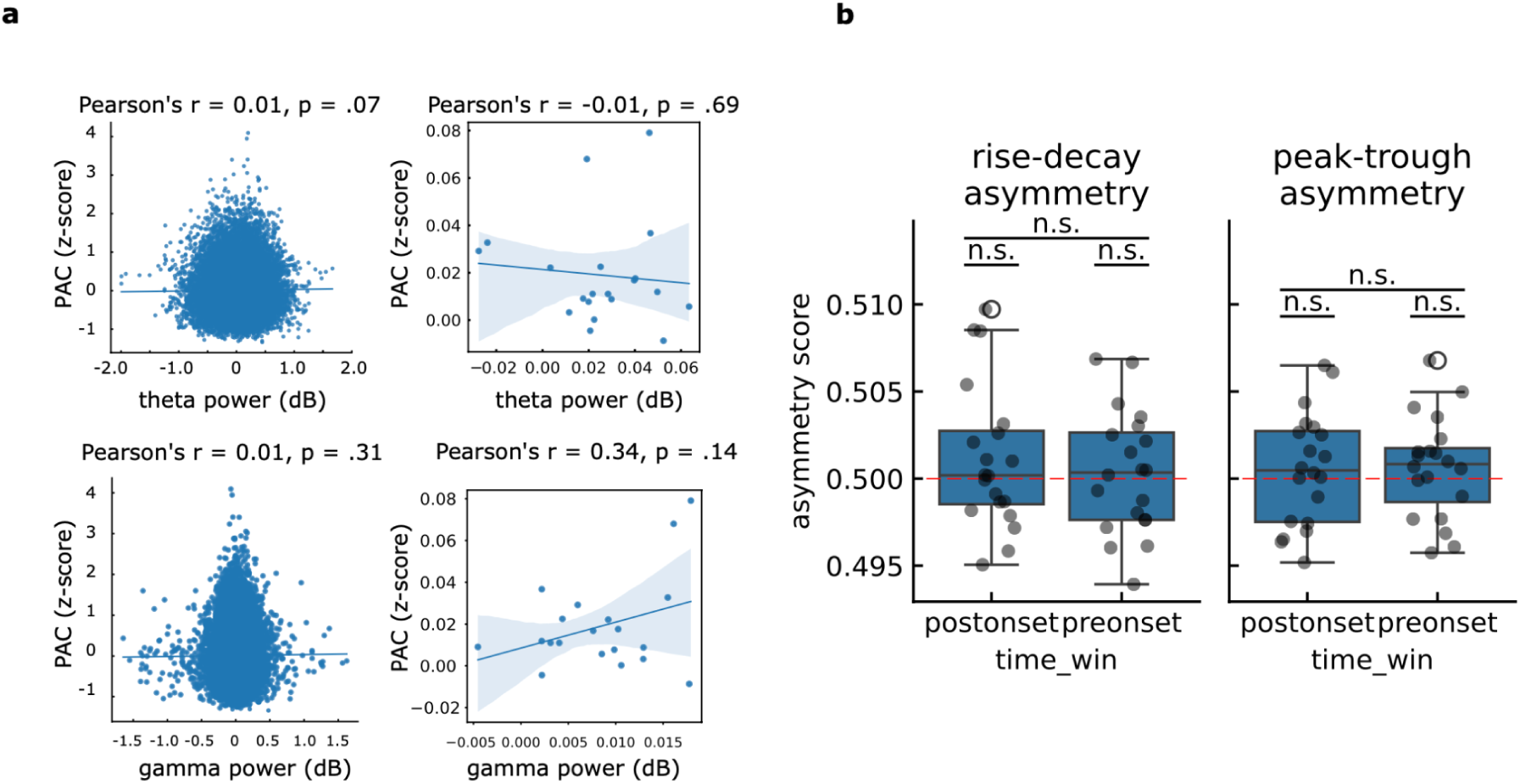
Control analyses for hippocampal theta-gamma PAC (Figure 4a of main text). (a) Hippocampal TG-PAC was not statistically significantly correlated with theta (upper row) or gamma (lower row) power, nor at individual trial level (left column) nor at single subject level (right column). (b) We did not observe any asymmetry in theta-waveform shape, which could lead to spurious TG-PAC (Cole & Voytek, 2017). Rise-decay asymmetry (left) was not statistically different than the expected value for a perfect sinusoid (0.5) neither in the preonset (t(19)=0.405, p=0.689) nor postonset (t(19)=1.234, p=0.231) time window, nor there was a statistically significant difference between the two (t(19)=-1.631, p=0.119). Similarly, peak-through asymmetry (right) was not statistically different than the expected value for a perfect sinusoid (0.5) neither in the preonset (t(19)=0.978, p=0.340) nor postonset (t(19)=0.765, p=0.453) time window, nor there was a statistically significant difference between the two (t(19)=0.239, p=0.813). Dots represent individual participants. Error bars represent standard error of the mean.

**Supplementary Figure 6.**
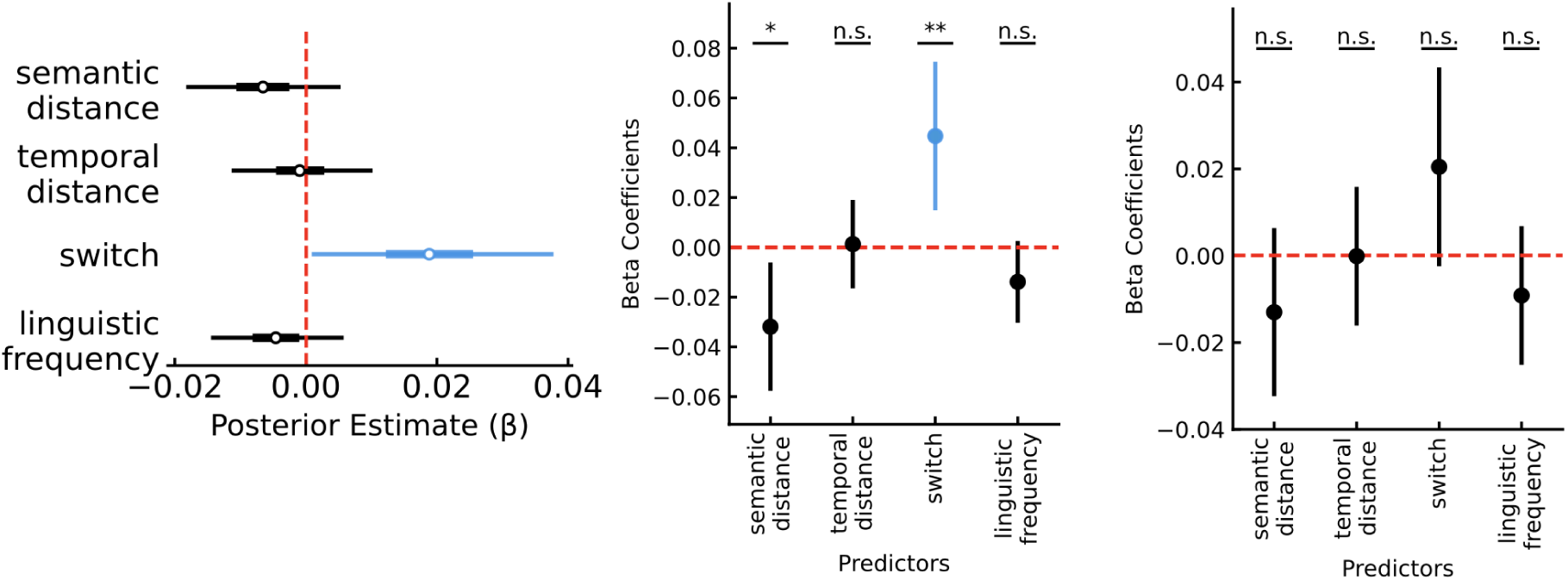
The effect reported in Figure 4b of main text (Bayesian full LMM) was mostly robust across different modelling approaches, namely Bayesian individual LMM (a): semantic distance (β = −0.007, 94% HDI = [−0.018, 0.005]), temporal distance (β = −0.001, 94% HDI = [−0.011, 0.01]), switch (β = 0.019, 94% HDI = [0.001, 0.038]), linguistic frequency (β = −0.005, 94% HDI = [−0.014, 0.006]); classical/frequentist full LMM (b): semantic distance (β = −0.031, CI= [−0.057, −0.006], p=0.027), temporal distance (β = 0.001, CI= [−0.016, 0.019], p=0.888), switch (β = 0.044, CI= [0.014, 0.074], p<0.001), linguistic frequency (β =-0.013, CI= [−0.030, 0.002], p=0.102); classical/frequentist individual LMM (c): semantic distance (β = −0.013, CI= [−0.032, 0.006], p=0.204), temporal distance (β = 0, CI= [−0.016, 0.015], p=0.990), switch (β = 0.020, CI= [−0.002, 0.043], p=0.083), linguistic frequency (β = −0.009, CI= [−0.025, 0.006], p=0.262). For the first two of them, we detected significant modulation of hippocampal TG-PAC as a function of switch transitions between subclusters.

**Supplementary Figure 7.**
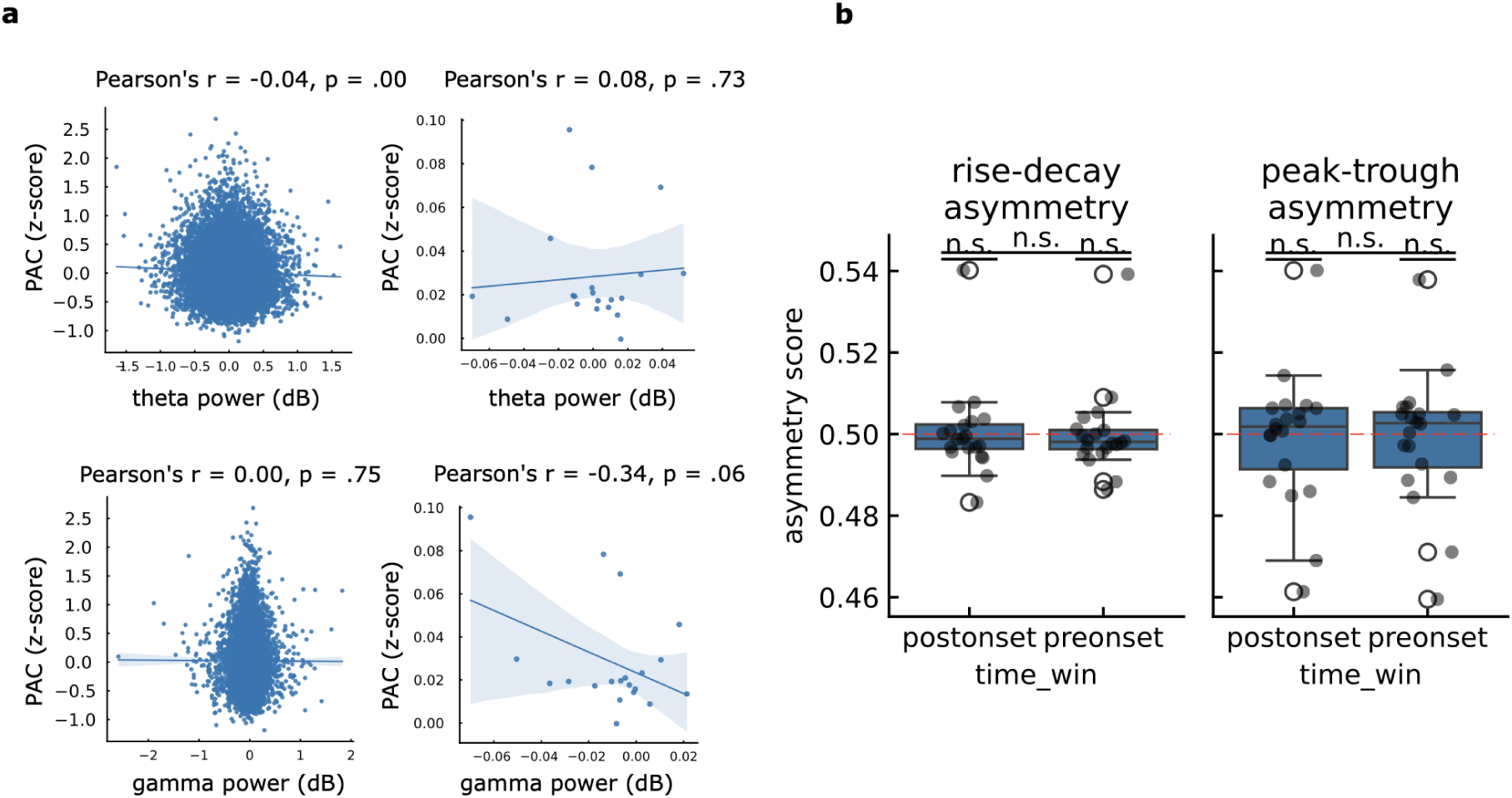
Control analyses for LTL theta-gamma PAC (Figure 4d of main text). (a) LTL TG-PAC was not statistically significantly correlated with gamma (lower row) power, nor at individual trial level (left) nor at single subject level (right). This was true also for theta power at single subject level (top right), but we did find a statistically significant effect when individual trials were considered. LTL TG-PAC therefore should be interpreted with caution, although the observed correlation score was rather modest (Pearson’s r = −0.04) (b) We did not observe any asymmetry in theta-waveform shape, which could lead to spurious TG-PAC (Cole & Voytek, 2017). was not influenced by theta waveform shape. Rise-decay asymmetry (left) was not statistically different than the expected value for a perfect sinusoid (0.5) neither in the preonset (t(19)=0.040, p=0.968) nor postonset (t(19)=0.134, p=0.894) time window, nor there was a statistically significant difference between the two (t(19)=-0.458, p=0.651). Similarly, peak-through asymmetry (right) was not statistically different than the expected value for a perfect sinusoid (0.5) neither in the preonset (t(19)=-0.290, p=0.774) nor postonset (t(19)=-0.286, p=0.777) time window, nor there was a statistically significant difference between the two (t(19)=0.003, p=0.997). Dots represent individual participants. Error bars represent standard error of the mean.

**Supplementary Figure 8.**
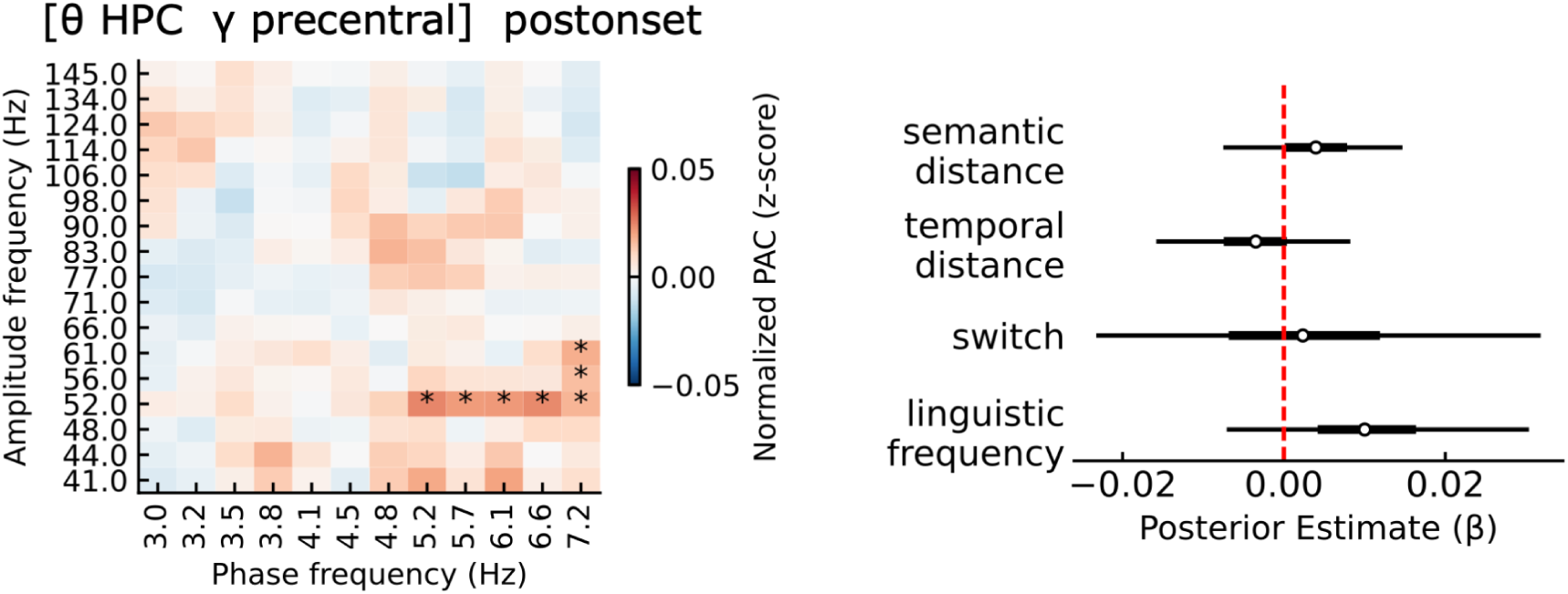
We observed a significant cross-regional TG-PAC between phase of hippocampal theta and amplitude of precentral gamma in the postonset period (left)(p < .05, cluster corrected). This, however, was not statistically significantly modulated by any of the behavioural factors we isolated (right): semantic distance (β = 0.004, 94% HDI = [−0.008, 0.015]), temporal distance (β = −0.004, 94% HDI = [−0.016, 0.008]), switch (β = 0.003, 94% HDI = [−0.023, 0.032]), linguistic frequency (β = 0.01, 94% HDI = [−0.007, 0.03]).

**Supplementary Figure 9.**
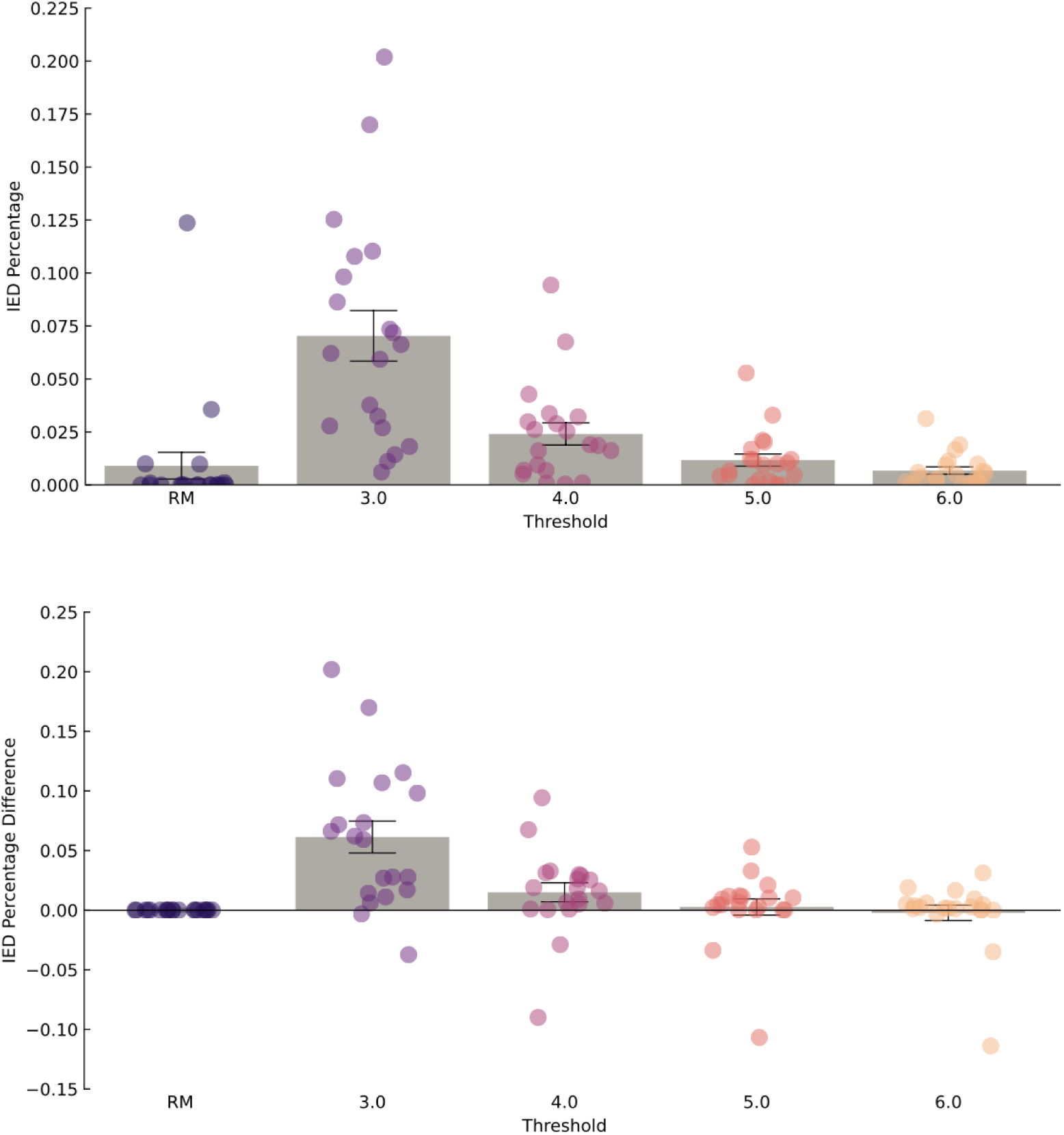
All recordings were visually inspected by an expert epileptologist (RM), who identified and marked electrode contacts and time windows contaminated by potential interictal epileptiform discharges (IEDs) for exclusion. To validate this clinical assessment, we compared the proportion of data excluded against that identified by an automated detection method (Gelinas et al., 2016). The results show that both approaches yield comparable exclusion rates, particularly at higher detection thresholds. The largest point of disagreement, with threshold of 3.0, resulted in a discrepancy of about 5%, which we interpreted as negligible. Increasing the threshold similarly to what has been reported in more recent work on spatial navigation (e.g., Chen, Kunz et al., 2018; Aghajan et al., 2017; Stangl et al., 2020; Maoz et al., 2023; Liu et al., 2023) resulted in an even smaller discrepancy. Given the absence of clear consensus on how to select the parameters of such automated detection methods, we thus relied on clinical expertise. The upper panel illustrates the percentage of IEDs flagged by the clinical assessment (RM) and by the automated method across varying thresholds. The lower panel depicts the difference between the two methods, relative to clinical assessment. Dots represent individual participants. Error bars represent standard error of the mean.

**Supplementary Table 1.** Descriptive data of participants, with sex, age in years, total number of trials included in the analyses (see Methods), contacts in hippocampus (Hip), LTL, and precentral, and hemisphere (L = left, R = right).

| Patient ID | Sex | Age (years) | # Trials included | Hip contacts | LTL contacts | precentral contacts | Hemisphere |
| --- | --- | --- | --- | --- | --- | --- | --- |
| 01 | M | 24 | 76 | 3 | 3 | 2 | L |
| 03 | F | 24 | 134 | 6 | 1 | 1 | L |
| 04 | F | 42 | 61 | 7 | 2 |  | L |
| 05 | F | 32 | 123 | 3 | 1 | 3 | L |
| 06 | F | 44 | 130 | 8 | 2 | 6 | L |
| 07 | F | 46 | 132 | 7 | 9 |  | L |
| 08 | M | 27 | 118 | 6 | 4 | 4 | R |
| 09 | M | 30 | 123 | 7 | 3 | 3 | L |
| 10 | M | 35 | 106 | 9 | 5 |  | L |
| 11 | M | 39 | 138 | 18 | 5 |  | L+R |
| 12 | M | 29 | 247 | 10 | 3 | 1 | R |
| 13 | M | 28 | 174 | 13 | 10 | 4 | L+R |
| 14 | M | 30 | 126 | 5 | 4 | 1 | R |
| 15 | F | 27 | 136 | 18 | 7 |  | L+R |
| 16 | M | 29 | 255 | 8 | 3 |  | L |
| 17 | F | 20 | 176 | 13 | 7 |  | L+R |
| 18 | F | 23 | 176 | 7 | 8 |  | L |
| 19 | F | 28 | 253 | 4 | 1 | 1 | R |
| 20 | F | 36 | 217 | 7 | 5 |  | R |
| 21 | M | 34 | 123 | 15 | 1 | 5 | R |
|  |  |  |  | 174 | 84 | 31 |  |

